# Longitudinal profiling of tumor-reactive T cells during TIL therapy in metastatic melanoma

**DOI:** 10.1101/2025.09.23.678066

**Authors:** Michael T. Sandholzer, Clara Serger, Alessia G. Liner, Sarp Uzun, David König, Helen Thut, Reto Ritschard, Jonas D. Fürst, Andreas Zingg, Natalia Rodrigues Mantuano, Benjamin Kasenda, Katharina Glatz, Elisabeth Kappos, Matthias Matter, Andreas Holbro, Frank Stenner, Jakob Passweg, Nina Khanna, Lukas Jeker, Mascha Binder, Alfred Zippelius, Heinz Läubli

**Affiliations:** Department of Biomedicine, University of Basel and University Hospital Basel, Switzerland; Division of Medical Oncology, University Hospital Basel, Switzerland; Department of Pathology, University Hospital Basel, Switzerland; Departement of Plastic, Reconstructive, Aesthetic and Hand surgery, University Hospital Basel and University of Basel; Division of Dermatology, University Hospital Basel, Switzerland; Division of Hematology, University Hospital Basel, Switzerland; Division of Infectious Diseases, University Hospital Basel, Switzerland; Innovation Focus Cell Therapies, University Hospital Basel, Switzerland

**Keywords:** cancer immunotherapy, T cell receptor, cell therapy, tumor-reactive T cells, melanoma

## Abstract

Adoptive cell therapy (ACT) with expanded autologous tumor-infiltrating lymphocytes (TILs) can induce durable responses in metastatic melanoma, yet many patients relapse. We profiled tumor-reactive T cell dynamics during TIL therapy using single-cell RNA and TCR sequencing from seven patients to elucidate underlying reasons. We found that tumor-reactive T cells preferentially expanded early during *ex vivo* TIL culture, transitioning from exhausted to reinvigorated effector states. Particularly, CD8^+^ exhausted T cells (Tex) and CD4^+^ follicular helper T cell (Tfh), but not CD4^+^ Tex, were efficiently reinvigorated. Further, we resolved the heterogeneity of tumor-reactive CD8^+^ and CD4^+^ subsets, defining unique signatures for their identification during TIL expansion. In addition, non-responders (NRs) exhibit increased levels of Type 17 T cells in TIL products, suggesting a potential association with resistance to therapy. After transfer, tumor-reactive clones rapidly extravasated and established a stem-like reservoir. However, in NRs, CD4^+^ regulatory T cells (Tregs) expanded *de novo* and tumor-reactive CD8^+^ T cells reacquired exhaustion markers, limiting their functionality. By contrast, responders (Rs) retained a pool of less differentiated, stem-like cells.

Collectively, these data provide a comprehensive analysis of T cell fates during TIL-ACT providing the basis for new approaches to enhance therapeutic strategies.

## Introduction

Metastatic melanoma remains one of the most challenging malignancies to treat, given its aggressiveness, capacity of immune evasion, and marked intra- and intertumoral heterogeneity. Despite advances in immune-checkpoint blockade (ICB) and targeted therapies such as BRAF and MEK inhibitors, durable responses are achieved only in a subset of patients, highlighting the need for novel therapeutic approaches^1,2^. Adoptive cell therapy (ACT) using tumor-infiltrating lymphocytes (TILs) has emerged as a promising treatment for metastatic melanoma. This approach involves isolating autologous T cells from tumor lesions, expanding them *ex vivo*, and re-infusing them into patients to enhance anti-tumor immunity, yielding objective response rates of up to 50% across multiple clinical trials in heavily pre-treated patients^3–5^. In a recent phase III trial, TIL-ACT more than doubled the progression-free survival of patients with advanced melanoma compared to ICB treatment^6^. This and other successful trials led to the FDA’s first approval of a commercial TIL therapy, authorized for patients’ refractory to standard therapies^7^. However, despite these advances, only a subset of patients derives long lasting clinical benefit from TIL therapy. Over the past decades, significant efforts have focused on elucidating the cellular and molecular mechanisms governing ACT responses. A critical determinant of ACT efficacy is the ability of transferred T cells to persist *in vivo*^8,9^. Studies have linked T cell persistence and anti-tumor efficacy to a less differentiated T cell phenotype with stem-like features, whereas fully differentiated effector T cells exhibit reduced persistence and impaired anti-tumor efficacy upon infusion^10–12^. While ACT approaches using engineered T cell receptors (TCRs) or chimeric antigen receptors (CARs) address this by transferring less differentiated cells, TIL-ACT offers a distinct and complementary advantage by leveraging the natural polyclonality of tumor-infiltrating T cells, enabling broad tumor recognition, even in the face of tumor antigen heterogeneity^13–15^. However, during the *ex vivo* expansion, tumor-dominant T cell clones are often outcompeted by newly emerging clones, reflecting complex and poorly understood T cell dynamics during TIL expansion^16^. To achieve the full therapeutic potential of TIL-ACT, it is critical to elucidate the mechanisms governing clonal dynamics, T cell persistence, and effector function.

In this study, we address these critical gaps by dissecting the dynamics and persistence of tumor-reactive T cell subpopulations during TIL therapy. Leveraging longitudinal samples from patients with advanced metastatic melanoma enrolled in the BaseTIL trial, we tracked tumor-reactive T cell clones throughout TIL therapy, examining their fate during *ex vivo* expansion, post-transfer in the peripheral blood, and in post-treatment metastatic lesions.

## Results

### Plasticity and clonal dynamics of tumor-reactive CD8^+^ T cells

To investigate T cell fate decisions during TIL-ACT, we collected longitudinal samples from seven patients participating in the phase I BaseTIL trial (NCT04165967), which evaluated TIL therapy in combination with the PD-1 inhibitor nivolumab for the treatment of metastatic melanoma^17^. Our cohort comprised two patients achieving partial response according to the Response Evaluation In Solid Tumors (RECIST) version 1.1, classified as responders (Rs), and five non-responders (NRs), including two patients with progressive disease, and three patients with stable disease. Patient samples were collected at five time points throughout TIL-ACT for longitudinal analysis (Fig. 1a). Baseline tumor sample that served as the TIL source (pre-ACT) (1). The following e*x vivo* TIL expansion comprises a two-step expansion protocol^18^: In the first step, tumor fragments are cultured in high-dose IL-2 to generate an intermediate product (2) enriched for tumor-infiltrating T cells, a process referred to as the pre-rapid expansion protocol (preREP). In the second step, the intermediate product undergoes a rapid expansion protocol (REP) using high-dose IL-2, anti-CD3 antibodies and allogeneic feeder cells to achieve sufficient cell numbers for therapeutic infusion, yielding the final TIL product (3). After *ex vivo* expansion, TILs are infused into lymphodepleted patients, accompanied by IL-2 administration to support TIL engraftment and persistence. After transfer we collected peripheral blood mononuclear cells (PBMC) 7 days post transfer (7dpt) (4), and from five patients metastatic lesions post transfer (post-ACT) (5) (Extended Data Table 1).

**Fig. 1:**
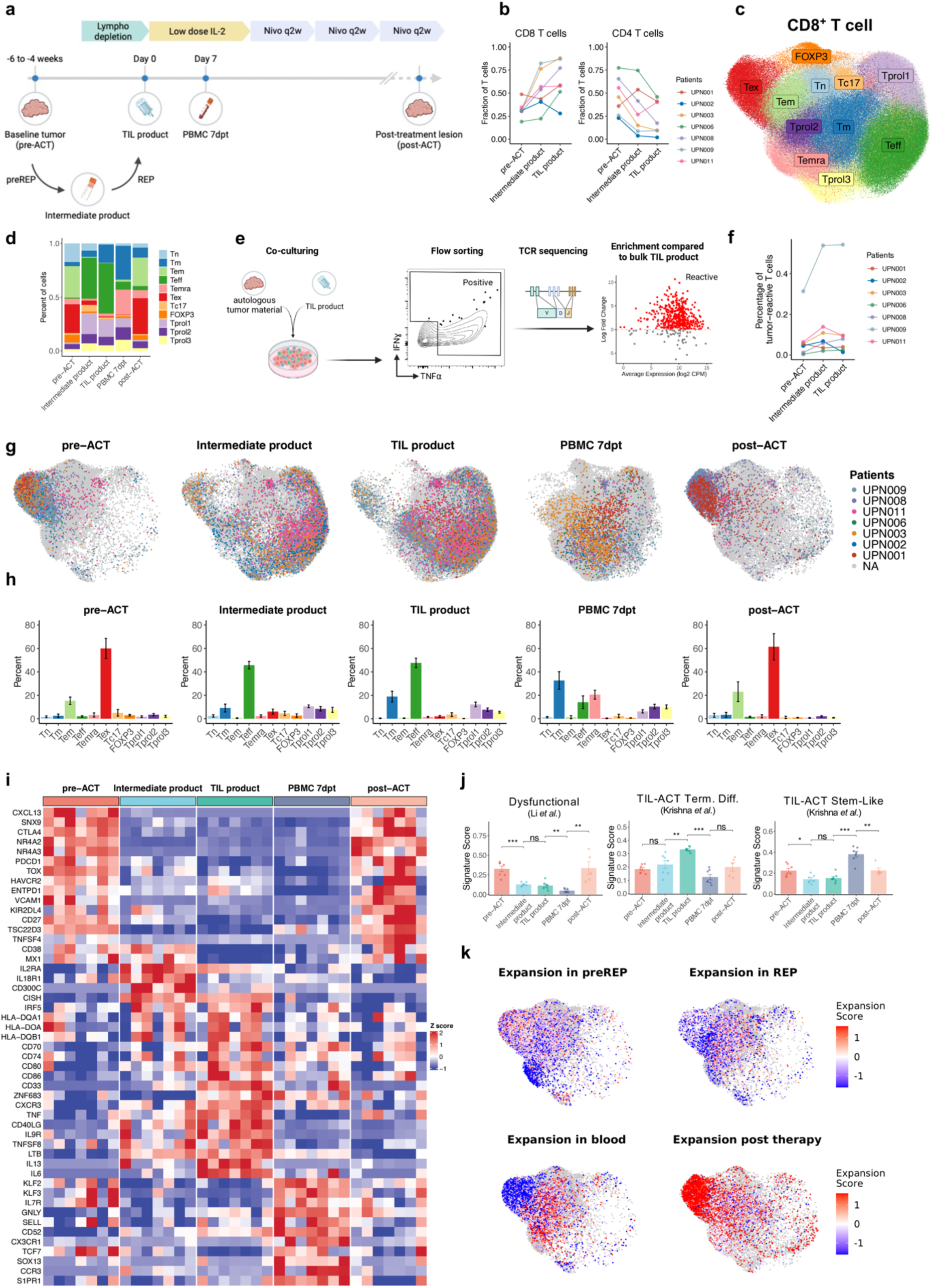
Plasticity and clonal dynamics of tumor-reactive CD8^+^ T cells. **a)** Treatment outline and time points of sample collection. Baseline tumor (pre-ACT, n=7); intermediate product (n=7); TIL product (n=7); peripheral blood mononuclear cells 7 days post transfer (PBMC 7dpt, n=7); post-treatment lesion (post-ACT, n=5+4); REP, rapid expansion protocol. Illustration was prepared using BioRender.com. **b)** CD8^+^ and CD4^+^ T cell frequencies during TIL expansion. **c)** UMAP visualization of CD8^+^ T cells. Clusters were annotated using their functional annotation or representative genes: Tn, naïve T cells; Tm, memory T cells; Tem, effector memory T cells; Teff, activated effector T cells; Temra, terminally differentiated effector memory or effector T cells; Tex, exhausted T cells; Tc17, Type 17 CD8^+^ T cells; FOXP3, CD8^+^ T cells expressing FOXP3; Tprol, T cells with high expression of proliferation markers. **d)** Proportional changes of CD8^+^ T cell subtypes during TIL therapy. **e)** Graphical summary of tumor-reactive T cell identification. Illustration prepared using BioRender.com. **f)** Percent of tumor-reactive T cells during TIL expansion. **g)** Tumor-reactive CD8^+^ T cells highlighted in the UMAP representation at collected time points. **h)** Bar plots showing the mean CD8^+^ T cell composition of the tumor-reactive subpopulation per time point. Error bars indicate SEM. **i)** Heatmap of the top differentially expressed genes in tumor-reactive CD8^+^ T cells between time points. **j)** Signature scores of patient bulked tumor-reactive CD8^+^ T cells during TIL expansion and post-transfer^12,26^. Statistical significance was assessed using Wilcoxon rank sum test. * p ≤ 0.05; ** p≤0.01; *** p≤0.001. **k)** Expansion score highlighted in the UMAP representation of pre-ACT or post-ACT.

Isolated T cells from these samples were subjected to single-cell RNA-sequencing (scRNA-seq) and TCR-sequencing (scTCR-seq). After filtering, high-quality expression data was obtained from 353’728 T cells, with TCRβ CDR3 regions in 284’339 cells (Extended Data Table 2 & 3). After dimensionality reduction, batch correction, and Leiden clustering, CD3^+^ T cells were separated into CD4^+^ and CD8^+^ subsets. During TIL expansion protocol, CD8^+^ T cells preferably expanded while CD4 T cells account for a reduced fraction in TIL products (Fig. 1b). The Uniform Manifold Approximation and Projection (UMAP) of CD8^+^ T cells revealed eleven distinct subpopulations that were annotated based on previously published transcriptional signatures^19^ and highly expressed marker genes (Fig. 1c and Extended Data Fig. 1a, b).

Given the high clonal overlap between timepoints, cluster analysis indicated substantial shifts in T cell populations during TIL therapy, with exhausted (Tex) and effector-memory (Tem) phenotype in pre-ACT converging to effector (Teff) and proliferating (Tprol) profiles during TIL expansion (Fig. 1d, Extended Data Fig. 1c). During this, the naïve T cell (Tn) population diminished, indicating T cell activation and differentiation during the REP phase. Seven days post transfer, during IL-2 administration, the proportion of Teff cells was markedly reduced and the majority of T cells displayed Tprol, Tm or terminally differentiated effector memory (Temra) profiles suggesting retained proliferation but reduced activation after transfer. Upon reencounter of the TME in tumor lesions collected post-ACT, the distribution of T cell profiles resembled that of pre-ACT baseline samples, with prominent Tex and Tem populations (Fig. 1d).

As tumor-reactive clones are critical for tumor control, we next focused on identifying these TCRs and tracing their fate in more detail throughout TIL therapy. Thus, we developed a tumor reactivity assay in which TIL products were co-cultured with autologous tumor-derived single-cell suspensions or short-term cultured primary melanoma cell lines (Fig. 1e and Extended Data Fig. 1f, g). Subsequently, tumor-reactive clones were identified by comparing TCRβ sequences from sorted IFNγ⁺ or TNFα⁺ T cells to those from bulk TIL products, defining enrichment based on log-fold change. The proportion of identified tumor-reactive T cells in pre-ACT tumor samples varied across patients (Fig. 1f). However, during preREP expansion, tumor-reactive clones generally increased in proportion but either plateaued or declined during the REP phase (Extended Data Fig. 1f). Consistent with findings by Chiffelle *et al.*^20^, tumor-reactive T cell clones predominantly displayed a Tex profile in pre-ACT samples and were reinvigorated during TIL expansion (Fig. 1g, h). Already in preREP, they transitioned to an activated Teff state, which became even more pronounced after REP. In the final TIL product, most tumor-reactive T cells displayed Teff characteristics with a smaller subset showing Tm features. However, by day seven post transfer, the majority of transferred tumor-reactive T cells had transitioned to a Tm or Temra profile, with only a small fraction retaining Teff characteristics. Notably, in post-ACT lesions, the majority of tumor-reactive T cells reacquired Tex features, closely resembling their pre-ACT counterparts (Fig. 1g, h).

This transcriptional shifts of tumor-reactive CD8^+^ T cells during TIL therapy were further investigated using differential gene expression (DGE) analysis comparing tumor-reactive clones between timepoints. This confirmed that in pre-ACT and post-ACT samples, tumor-reactive T cells showed high expression of exhaustion-associated markers (*PDCD1, CTLA4, TOX, KIR2DL4*) (Fig. 1i). During preREP, this exhausted profile was downregulated, with upregulation of *IL2RA*, *HLA-DOA,* and *CD74,* genes linked to activation, antigen processing and presentation, as well as inhibitors of effector function and proliferation (*CISH, CD300C*)^21,22^ suggesting limited expansion potential (Fig. 1i). In the final TIL product, tumor-reactive clones showed elevated expression of co-stimulatory ligands (*CD80*, *CD86*), activation markers (*TNF, CD40LG*), and genes involved in tumor homing and tissue-residency (*CXCR3, ZNF683*)^23^ (Fig. 1i). Notably, by seven days post transfer, we observed the same set of clones with an increased expression of markers associated with stemness (*TCF7, SELL, IL7R*) and immune cell trafficking (*KLF2*, *S1PR1, SELL, CX3CR1)*, suggesting active trafficking through lymphoid tissue^24,25^ (Fig. 1i). These shifts in transcriptional profiles were further confirmed using publicly available gene signatures^12,26^, highlighting plasticity from dysfunctional to terminal differentiated profiles during TIL expansion but stem-like features after transfer (Fig. 1j).

This data revealed considerable T cell adaption to different environments including T cell activation and proliferation but also inhibition throughout different steps of TIL therapy. Hence, in a next step we calculated expansion (proportional increase) or contraction (proportional decrease) of clones across consecutive samples to elucidate clonal dynamics on single cell level (Fig. 1k). This analysis revealed that during preREP, especially tumor-reactive T cells in Tex cluster expanded significantly, indicating an IL-2-induced expansion of already activated T cells. During the REP phase, clone-independent stimulation by anti-CD3 antibodies caused robust expansion of Tn and Tem clones while tumor-reactive clones plateau or contracted indicating competitive disadvantage. Consistent with this, we observed a decline in the number of unique T cell clones (TCR richness) during the preREP phase, followed by an increase during the REP phase, suggesting that initially, only a subset of clones underwent expansion (Extended Data Fig. 1d). By day seven post-transfer, Tem and Temra clones dominated peripheral blood, with fewer tumor-reactive clones than in the TIL product (Fig. 1k). This finding aligns with the observed acquisition of stemness and immune cell trafficking markers in peripheral blood and suggest that these clones rapidly exit the circulation, infiltrate tissues, and later reappear in peripheral blood after migrating through lymph nodes. To support this assumption, we evaluated tumor infiltration and expansion of the transferred clones by comparing proportional changes between PBMCs at 7dpt and post-ACT lesions (Fig. 1k). Notably, especially tumor-reactive clones infiltrated post-ACT lesions and were found highly expanded in a Tex state.

Taken together, these data demonstrate that tumor-reactive CD8^+^ T cells undergo substantial clonal and transcriptional alterations during *ex vivo* expansion and after reinfusion into the patient. *Ex vivo* expansion reinvigorated tumor-reactive CD8^+^ TILs, shifting their transcriptional profiles from a canonical exhausted state toward an activated effector state, already during preREP. Notably, tumor-reactive CD8⁺ T cells expand during preREP but are subsequently outcompeted by bystander T cells during REP. After transfer, these tumor-reactive clones show increased immune trafficking potential and display a stem-like profile in the peripheral blood, suggesting newly generated tumor-reactive T cell in the LN niche^25^. Even though tumor-reactive CD8^+^ T cells successfully infiltrated and expanded in post-ACT lesions, they reacquire exhaustion features, highlighting the immunosuppressive effects of the TME.

### Tumor-reactive CD8^+^ T cells acquire distinct expression profiles during reinvigoration

Because tumor-reactive T cells had already expanded and been functionally reinvigorated during preREP, yet were often outcompeted during REP, we next examined their expression profiles in the intermediate products in order to identify unique features to develop strategies to improve TIL expansion.

First, we performed DGE comparing tumor-reactive CD8^+^ T cells with non-tumor reactive populations to resolve their unique expression profile (Fig. 2a). Tumor-reactive CD8^+^ T cells were mainly characterized by high expression of genes associated with antigen processing and presentation (*HLA-DQA1, HLA-DOA, CD74*), as well as co-costimulatory ligands (*CD80*, *CD86, CD70*). However, while retaining expression of co-inhibitory receptors (*PDCD1*, *CTLA4*) and inhibitors of T cell proliferation (*CD300C*)^21^, they lack expression of stemness features (*TCF7*, *IL7R*, *LEF1*, *CCR7*), which may explain their proliferative disadvantage compared to naive bystander populations during REP. Building on these observations, we next subclustered intermediate products of 6 patients revealing seven distinct clusters (Fig. 2b). Highlighting the identified tumor-reactive clones showed that they primarily populated cluster 1 and cluster 4 (Fig. 2c, Extended Data Fig. 2a). Moreover, clusters 1 and 4 showed high clonal overlap, indicating clone-independent development, where the same clone can adopt both states (Fig. 2d). Since cluster 1 is characterized by high expression of genes known for HLA-II antigen presentation and low levels of exhaustion markers, it may represent cells early after reinvigoration (Fig. 2e). In contrast, cluster 4 is characterized by *KLF2* expression, which is inversely regulated by TCR signaling^27^, and genes often linked to immune cell trafficking and migration (*S1PR1*, *SELL*, *CX3CR1*, *CXCR1*), suggesting a subset of tumor-reactive T cells not recently engaged in antigen recognition (Fig. 2e).

**Fig. 2:**
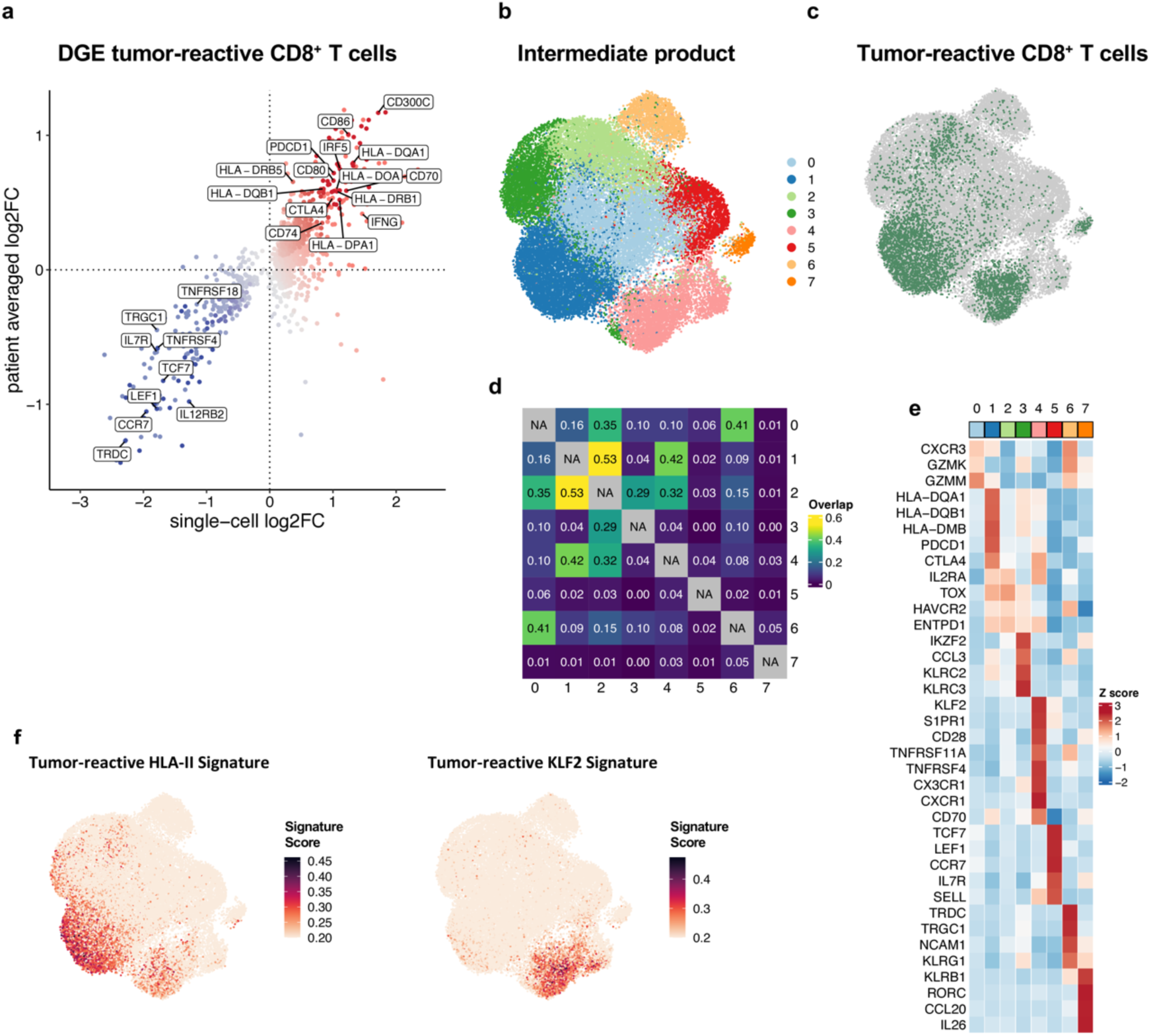
Tumor-reactive CD8^+^ T cells acquire distinct expression profiles during reinvigoration. **a)** Differential gene expression comparing tumor-reactive CD8^+^ subpopulations with bystander T cells in intermediate products using patient bulked comparison and single-cell comparison. Genes significantly upregulated in both comparisons were highlighted in dark color and genes significantly upregulated only in single-cell comparison in light color. **b)** UMAP visualization of CD8^+^ T cell states in intermediate products. **c)** Tumor-reactive CD8^+^ T cells highlighted in UMAP visualization of intermediate products. **d)** Heatmap presenting clonal overlap between clusters. **e)** Top expressed genes per cluster. **f)** Tumor-reactive HLA-II and Tumor-reactive KLF2 Signature highlighted on UMAP.

Finally, we defined subtype-specific signatures to assess whether the two states represent distinct transcriptional profiles (Fig 2f and Extended Data Fig. 2b, c). Clusters 1 and 4 both score high for the tumor-reactive HLA-II signature, but only cluster 4 also scores high for the tumor-reactive KLF2 signature, suggesting a transition from an HLA-II high to a KLF2 high state.

In conclusion, these data suggest that tumor-reactive T cells with an exhausted profile in pre-ACT tumors downregulate exhaustion markers during IL-2 exposure in preREP, developing toward an HLA-II high state. If no continuous TCR stimulus is provided, they may further transition into an KLF2 high state exhibiting features of increased immune trafficking. Notably, both states can be defined by unique signatures allowing quantification or isolation during TIL expansion protocols.

### Tumor-reactive CD8^+^ T cells exhibit dysfunctional profile in post-ACT of NRs

We next sought to determine whether *ex vivo* expansion and transfer had a profound impact on tumor-reactive T cell expression profiles after therapy and if reacquisition of exhaustion in tumor-reactive T cells correlates with therapeutic outcome.

Therefore, we compared CD8^+^ tumor-reactive cells from pre-ACT lesions (n=6) and post-ACT lesions (n=5) from six patients, including two Rs and four NRs. Unsupervised clustering of 9’062 tumor-reactive CD8^+^ T cells revealed six distinct clusters (Fig. 3a and Extended Data Fig. 3a, b). To distinguish late from early T cell differentiation states, we performed trajectory analysis (Fig. 3b and Extended Data Fig. 3e, f). This revealed two distinct paths originating from Tn and progenitor exhausted (Tpex) states: One trajectory led to ITGAE^+^ Tex and subsequently to CXCL13^+^ Tex cells. The second path led toward GZMK^+^ Tex and progressed to terminally exhausted T (Ttex) cells (Fig. 3b and Extended Data Fig. 3e, f). Notably, comparing pre- and post-ACT lesions, NRs showed enrichment of terminally differentiated CXCL13⁺ Tex and Ttex with reduced Tn and Tpex, whereas Rs exhibited increased GZMK⁺ Tex, Tpex, and Tn populations (Fig. 3c). Gene set enrichment analysis (GSEA) on matched clones from pre- and post-ACT confirmed this observation on a single clone level (Fig. 3d, f). In NRs, transferred tumor-reactive clones showed increased expression of dysfunctional^26^ and terminally differentiated^28^ gene signatures, accompanied by a loss of stem-like features^28^ post-ACT (Fig. 3d and Extended Data Fig. 3g). In contrast, clones of a R retained stem-like T cell signatures with reduced dysfunctional^26^ and terminally differentiated^28^ profiles previously associated with favorable responses in immunotherapy^29^ (Fig. 3f and Extended Data Fig. 3g). To further dissect transcriptional changes in tumor-reactive clones between pre-ACT and post-ACT samples of Rs and NRs, we performed DGE analysis on the top expanded clones (Fig. 3e). While tumor-reactive clones in pre-ACT showed expression of markers for infiltration, tissue-residency and early activation (*VCAM1, ITGAE, NR4A3, NFKB1, IRF1*), clones of R post-ACT mainly upregulated granzymes (*GZMK*, *GZMA*, *GZMM*, *GHMH*) indicating cytotoxicity. In contrast, clones of NRs post-ACT exhibited high expression of inhibitory receptors and exhaustion markers (*PDCD1, CTLA4, HAVCR2*) suggesting a dysfunctional state (Fig. 3e).

**Fig. 3:**
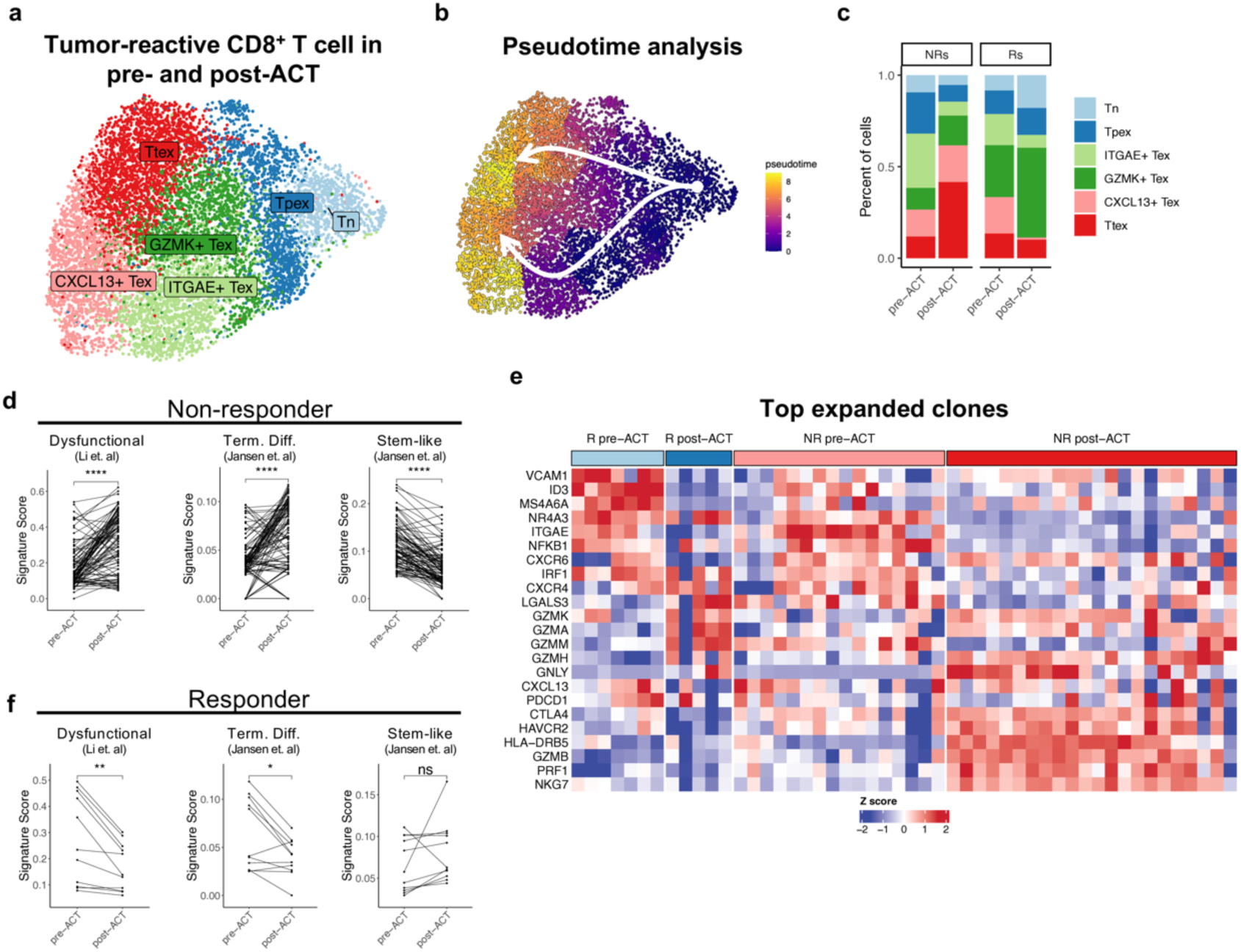
CD8^+^ tumor-reactive T cells show an elevated dysfunctional profile in post-ACT lesions of non-responders. **a)** UMAP visualization of tumor-reactive CD8^+^ T cell states in pre- and post-ACT lesions. Tn, naïve T cells; Tex, exhausted T cells; Tpex, precursors exhausted T cells; Ttex, terminally exhausted T cells. **b)** Pseudotime trajectories visualized on UMAP. **c)** Cluster distribution of tumor-reactive CD8^+^ T cells detected in pre- and post-ACT lesions of responders (Rs) and non-responders (NRs). **d, f)** Signature scores of published signatures comparing mapped clones of pre- and post-ACT lesions of Rs and NRs^26,28^. Statistical significance was assessed using Wilcoxon signed-rank test. * p ≤ 0.05; ** p≤0.01; *** p≤0.001; **** p≤0.0001. **e)** Heatmap showing the mean relative expression of differentially expressed genes in expanded clones (>15 cells) in pre- and post-treatment lesions of Rs and NRs.

Overall, these findings demonstrate that tumor-reactive CD8^+^ T cells in NRs are driven toward terminal exhaustion and dysfunction post-ACT. In contrast, responders retain a pool of less differentiated T cells with cytotoxic effector function. Importantly, these transcriptional changes are direct consequences of TIL therapy response and underscore the importance of preserving early-stage T cell phenotypes and highlight terminal differentiation as a key barrier to therapeutic efficacy.

### Plasticity and clonal dynamics of tumor-responsive CD4^+^ T cells

Having shown that tumor-reactive CD8^+^ T cells can undergo major transcriptional changes during TIL therapy, we next examined CD4^+^ T cells. Unsupervised clustering revealed nine distinct CD4**^+^** T cell subsets (Fig. 4a), which were annotated using signatures by Zheng *et al.*^19^ or identified by marker genes (Extended Data Fig. 4a, b). Similar to CD8^+^ T cells, CD4^+^ T cell subpopulations exhibited dynamic changes during TIL expansion and after transfer (Fig. 4b). In pre-ACT lesions, CD4**^+^** T cells primarily exhibited transcriptional characteristics of Tn, memory T cells (Tm), T helper 17 cells (Th17), and regulatory T cells (Treg) (Fig. 4b). During the pre-REP and REP phases, CD4**^+^** T cells transitioned primarily to a Teff phenotype, with increased fractions of Th17 and reduced Tregs. Seven days post transfer, we observed a strong increase in Tregs and a decrease in Teff proportions in the peripheral blood. Notably, T cell profiles in post-ACT lesions largely mirrored those from pre-ACT samples but displayed higher fractions of Treg, Teff and Th17 cells.

**Fig. 4:**
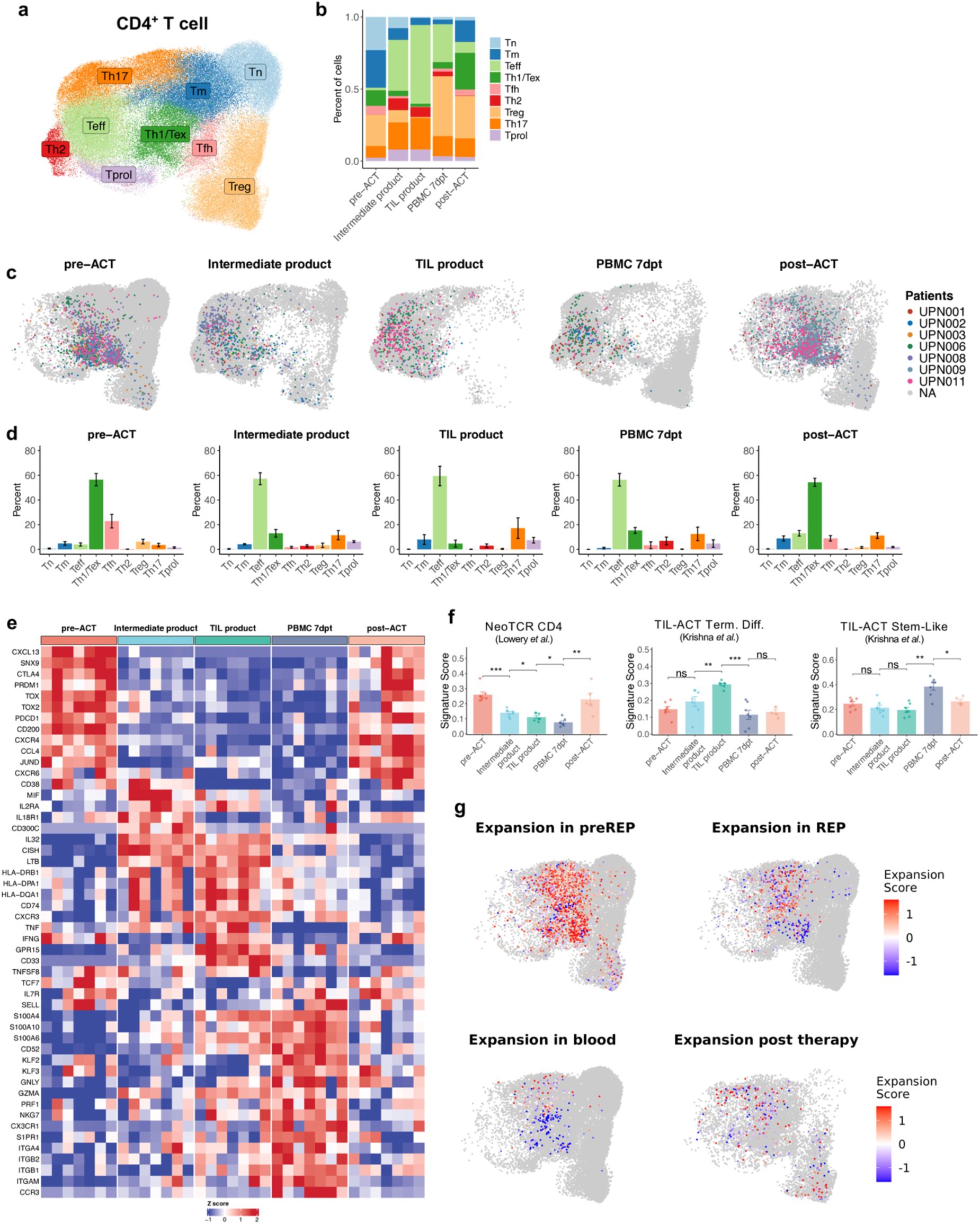
Plasticity and clonal dynamics of tumor-responsive CD4^+^ T cells. **a)** UMAP visualization of CD4^+^ T cell states. Clusters are highlighted using their functional annotation: Tm, memory T cells; Teff, activated effector T cells; Th1/Tex, T helper 1 cells and exhausted T cells; Tfh, T follicular helper cells; Th2, T helper 2 cells; Treg, regulatory T cells; Th17, T helper 17 cells; Tprol, T cells with high expression of proliferation markers. **b)** Proportional changes of CD4^+^ T cell subtypes during TIL therapy. **c)** Tumor-responsive CD4^+^ T cells highlighted in the UMAP representation at collected time points. **d)** Bar plots showing the mean CD4^+^ T cell composition of the tumor-responsive subpopulation per time point. Error bars indicate SEM. **e)** Heatmap of the top differentially expressed genes in tumor-responsive CD4^+^ T cells between time points. **f)** Signature scores of patient bulked tumor-responsive CD4^+^ T cells during TIL expansion and post-transfer^12,31^. Statistical significance was assessed using Wilcoxon rank sum test. * p ≤ 0.05; ** p≤0.01; *** p≤0.001. **g)** Expansion score highlighted in the UMAP representation of pre-ACT or post-ACT.

Although CD4⁺ T cells can exert direct cytotoxicity, only a minority of tumor-responsive clones are expected to do so, with most antitumor activity mediated through indirect mechanisms such as cytokine secretion, dendritic cell licensing, or support of CD8⁺ T cells^30^. To capture the broader landscape of tumor-responsive CD4⁺ T cells, we derived a transcriptional signature from experimentally validated tumor-reactive clones and applied it to conventional CD4⁺ T cells to identify additional clones with antitumor profiles resembling those directly tumor-reactive. Using this signature, which associated with CD4⁺ T cell expansion and aligned with established neoantigen-reactive CD4⁺ T cell signatures^31^ (Extended Data Fig. 4c-f), we identified tumor-responsive clones and monitored their transcriptional dynamics during TIL therapy.

This approach revealed that tumor-responsive clones in pre-ACT mainly exhibited a T helper 1/exhausted (Th1/Tex) and follicular helper T cell (Tfh) profile (Fig. 4c, d) and transitioned after preREP to a Teff profile, while only a fraction remained in a Th1/Tex state. During REP, nearly all tumor-responsive clones converged to a Teff state, while a small subset of Th17 expanded. Seven days after TIL transfer, most tumor-responsive clones in blood retained an activated Teff state, while some reverted to a Th1/Tex profile. Within post-ACT, the majority of tumor-responsive clones transitioned back to Th1/Tex, with only a fraction retaining the Teff state. However, compared to pre-ACT, more tumor-responsive clones acquired Teff, Th17 and Tm profiles, while fewer exhibited a Tfh state, suggesting a loss of tumor-responsive Tfh cells during TIL therapy.

Considering their notable plasticity throughout TIL therapy, we next performed DGE comparing the tumor-responsive CD4^+^ subsets between timepoints to further dissect changes in expression profiles. As expected, tumor-responsive CD4⁺ clones expressed high levels of anti-tumor and exhaustion markers (*CXCL13, PDCD1, TOX*) in pre-ACT, which were downregulated upon *ex vivo* expansion, suggesting reinvigoration. Similar to their CD8^+^ counterparts, in intermediate products, tumor-responsive CD4^+^ clones upregulated genes for HLA-II antigen presentation (*HLA-DRB1, CD74*) and markers for activated cytotoxic CD4^+^ T cells (*TNF, LTB, IL2RA*), but also inhibitors of T cell effector function and proliferation (*CISH, CD300C*)^21,22^. Some transcripts where even more pronounced in TIL products where high expression of *TNF, IFNG* and *CXCR3* suggests high activation and tumor homing potential (Fig. 4e). Seven days post-transfer in PBMCs, the same clones expressed markers of immune trafficking and migration (*KLF2, S1PR1, CX3CR1*), stemness (*TCF7, IL7R, SELL*), and cytotoxicity (*GNLY, GZMK, PRF1, NKG7*), indicating a stemness reservoir capable of both tissue homing and effector function (Fig. 4e). Finally, in post-ACT lesions, tumor-responsive clones show again markers of anti-tumor response (*CCL4, TNF, IFNG*), *CXCR6,* which is critical for sustained tumor control^32^, but also immune checkpoints (*PDCD1, CTLA4*) (Fig. 4e). GSEA of the tumor-responsive subpopulation using published signatures^12,31^ confirmed that anti-tumor-associated programs were largely absent during ex vivo expansion, where cells displayed terminal differentiation profiles, but re-emerged in peripheral blood with a stem-like expression signature (Fig. 4f).

Having outlined the phenotypic and transcriptional changes tumor-responsive CD4^+^ T cells undergo during TIL therapy, we next turned to the clonal level to investigate how distinct T cell subsets expand or contract throughout the process. Especially tumor-responsive Th1/Tex and Tfh subsets underwent significant expansion in the preREP phase but contracted in the REP phase (Fig. 4g). During this, also Th17 cells expanded in preREP and partially in REP phase explaining increased fraction in final TIL products (Fig. 4b). After transfer, similar to their CD8^+^ counterparts, also tumor-responsive CD4^+^ clones disappeared from circulation implying active tissue homing (Fig. 4g). However, contrary to our expectations in post-ACT mainly Th17 or Treg subpopulations infiltrated from circulation and expanded, underscoring loss of beneficial tumor-responsive CD4^+^ T cells during TIL therapy.

Taken together, these results show that *ex vivo* expansion markedly remodels the transcriptional profiles of tumor-responsive CD4^+^ clones, shifting them from an exhausted anti-tumoral profile toward cytotoxic and antigen-presenting profiles with enhanced potential for tumor homing. Similar to tumor-reactive CD8^+^ T cells, tumor-responsive CD4^+^ clones expanded efficiently during the first phase but became diluted in the second phase. Following adoptive transfer, these clones displayed stemness, tissue-homing, and cytotoxic potential, but showed limited tumor infiltration. Notably, Th17 subpopulations expanded during *ex vivo* expansion and infiltrated along with Tregs post-treatment lesions, potentially influencing the anti-tumor response.

### Tumor-responsive Tfh and Tex retain effector function in post-ACT lesions

To further explore TIL therapy-induced changes in tumor-responsive CD4⁺ T cells and their impact on the anti-tumor response, we subclustered them from pre-ACT (n=6) and post-ACT (n=5) lesions from six patients, including two Rs and four NRs (Fig. 5a and Extended Data Fig. 5a-c). In post-ACT, NRs showed fewer tumor-responsive Tfh cells and more cells transitioning into a Ttex state (Fig. 5b), whereas Rs displayed no major population changes. This suggests that loss of Tfh cells together with an increase in Ttex phenotype may be linked to resistance. Subsequent pseudotime analysis confirmed Ttex and subsets of Tex as the terminal populations, which also displayed the highest clonal expansion (Fig. 5c, d). Comparing pre- and post-ACT tumor-responsive CD4^+^ T cell distribution in NR, we observed an overall shift toward terminal subpopulations (Fig. 5e). Specifically, in pre-ACT samples, the majority of cells exhibit either a naïve or Tfh profile, whereas in post-ACT samples, these cells transitioned toward Th1 and Ttex states.

**Fig. 5:**
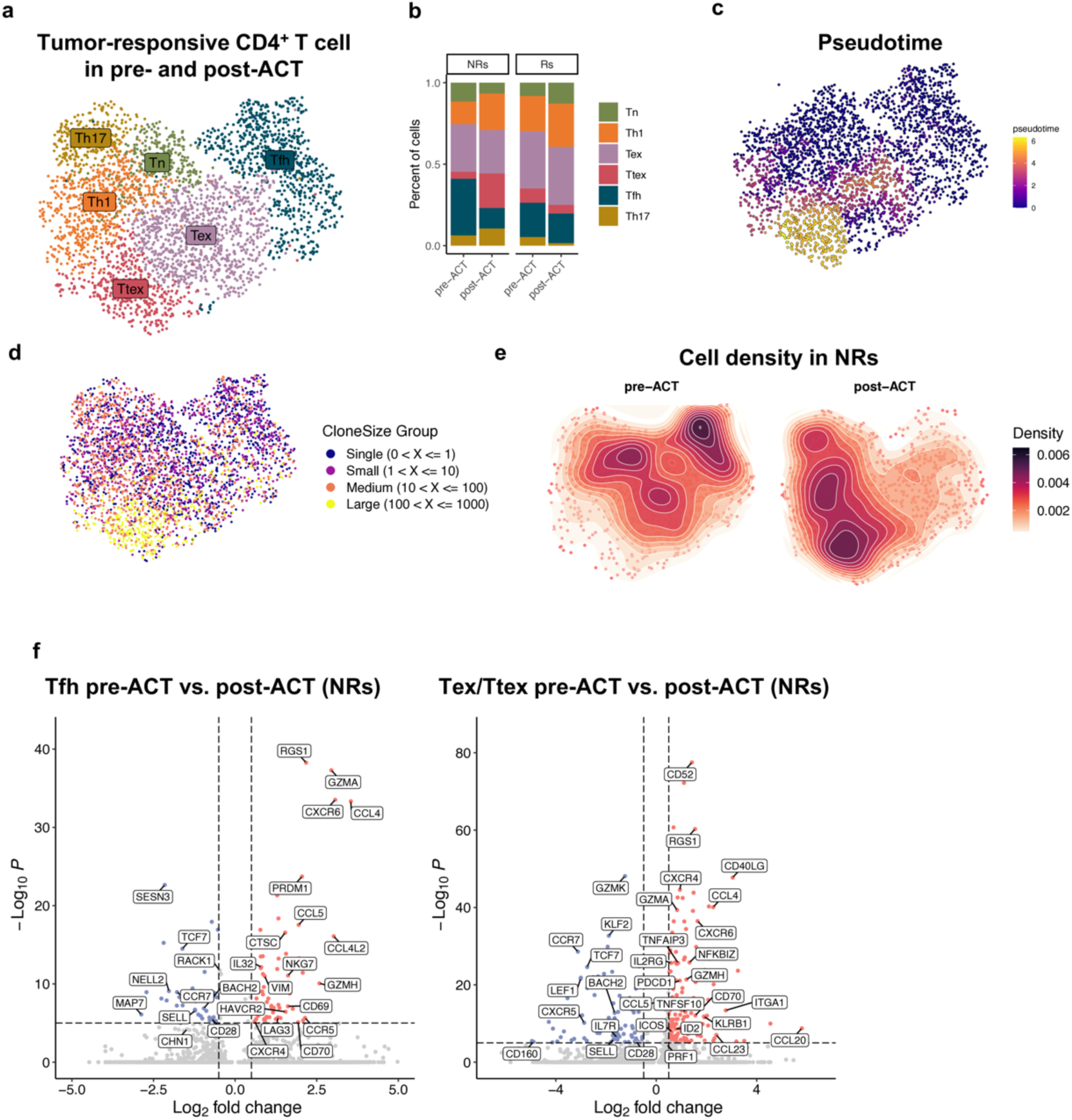
Tumor-responsive Tfh and Tex retain effector function despite immunosuppressive TME. **a)** UMAP visualization of tumor-responsive CD4^+^ T cell states in pre- and post-ACT lesions. Tn, naïve T cells; Th1, T helper 1 cells; Th17, T helper 17 cells; Tfh, T follicular helper cells; Tex, exhausted T cells **b)** Cluster distribution of CD4^+^ tumor-responsive T cells detected in pre- and post-ACT lesions of responders (Rs) and non-responders (NRs). **c)** Pseudotime scores visualized on UMAP. **d)** Expansion state highlighted in the UMAP representation. **e)** Cell density highlighted on UMAP of pre-ACT and post-ACT samples in NRs **f)** Volcano plot showing genes differentially expressed in tumor-responsive Tfh and Tex/Ttex comparing pre- vs. post-ACT in NRs.

To examine transcriptional changes in the main shifting populations, we performed DGE on Tfh and Tex/Ttex comparing pre- and post-ACT in NRs. In both populations, DGE analysis revealed a loss of stemness markers (*TCF7, SELL, CCR7*), an upregulation of Th1 markers (*CXCR6, PRDM1, TNFSF10*) and cytotoxicity-associated genes (*CCL4, CCL5, GZMA, GZMH*), indicating transition toward Th1-like effector states (Fig. 5f).

Taken together, both tumor-reactive CD8⁺ and CD4⁺ T cells in NRs expanded and progressed toward terminal, more mature states. While CD8⁺ T cells acquired a dysfunctional profile, CD4⁺ T cells instead gained enhanced effector function, albeit with a loss of stemness features.

### Tumor-responsive Tfh but not Tex were efficiently reinvigorated during preREP

Having established that tumor-responsive CD4⁺ T cells retain effector potential in post-ACT lesions, we next assessed whether the preREP expansion phase contributes to their reinvigoration. To address this, we compared tumor-responsive CD4⁺ T cells with other CD4⁺ T cells in intermediate products. Tumor-responsive clones expressed high levels of HLA-II antigen presentation genes (*HLA-DRB1, CD74*) and co-stimulatory ligands (*CD80, CD70*), but also showed exhaustion markers (*TOX, PDCD1*) and inhibitors of proliferation (*CD300C*)^22^, pointing to reinvigoration accompanied by partial dysfunction (Fig. 6a).

**Fig. 6:**
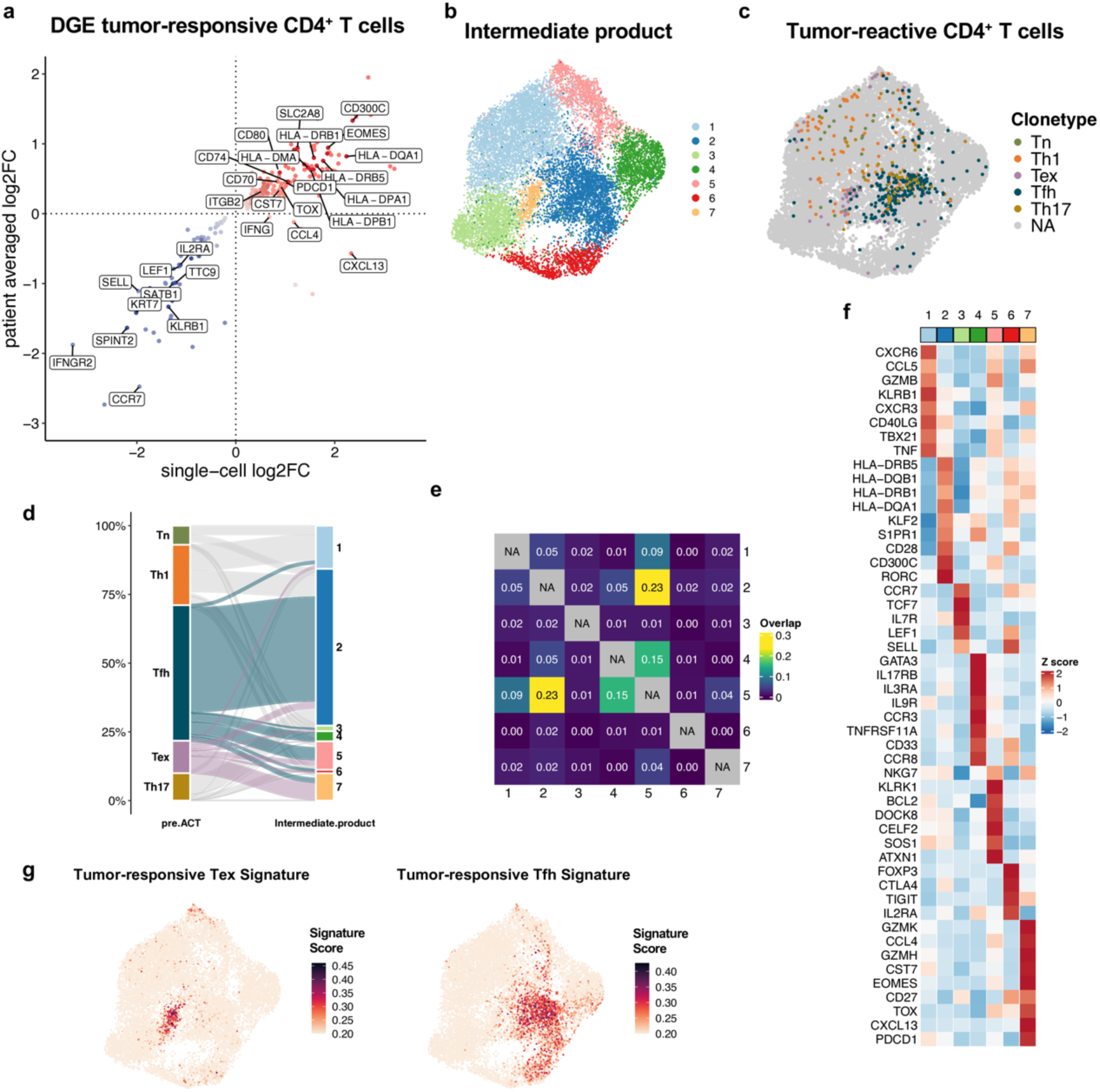
Tumor-responsive Tfh but not Tex were efficiently reinvigorated during preREP. **a)** Differential gene expression comparing tumor-responsive CD4^+^ subpopulations with bystander T cells in intermediate products using patient bulked comparison and single-cell comparison. Genes significantly upregulated in both comparisons were highlighted in dark color and genes significantly upregulated only in single-cell comparison in light color. **b)** UMAP visualization of CD4^+^ T cell states in intermediate products. **c)** Tumor-responsive CD4^+^ clone types highlighted in UMAP visualization of intermediate products. **d)** Alluvial plot depicting cluster localization of tumor-responsive clone types identified in pre-ACT on intermediate product UMAP. **e)** Heatmap presenting clonal overlap between clusters. **f)** Top expressed genes per cluster. **g)** Tumor-responsive Tex and Tumor-responsive Tfh Signature highlighted on UMAP.

To resolve which transcriptional profiles these clones adopt after preREP expansion, we subclustered intermediate products from six patients, identifying seven distinct CD4⁺ clusters (Fig. 6b). Mapping tumor-responsive clones revealed that Tfh-derived cells localized mainly in cluster 2, marked by enhanced HLA-II antigen presentation and migration (*KLF2, S1PR1*), resembling reinvigorated profiles similar to CD8⁺ T cells (Fig. 6c-f). In contrast, Tex-derived clones localized in cluster 7, characterized by cytotoxic activity (*CXCL13, GZMK, CCL4*) but persistent exhaustion (*PDCD1, TOX*), suggesting inefficient reinvigoration (Fig. 6f).

Notably, no clonal overlap was observed between clusters 2 and 7, indicating lineage-dependent rather than shared developmental trajectories during preREP expansion (Fig. 6e). Finally, we performed detailed DGE for Tfh and Tex subpopulations in intermediate products in order to identify unique signatures (Extended Data Fig, 6a, b). These signatures were then mapped onto the UMAP of intermediate products, confirming that they specifically highlighted reinvigorated Tfh and non-reinvigorated Tex populations, with no overlap (Fig. 6g).

Taken together, these data indicate that high-dose IL-2 during preREP selectively reinvigorates Tfh but not Tex clones, generating distinct transcriptional profiles. Importantly, the unique signatures we identified enable the isolation and further characterization of these subpopulations, potentially improving future therapeutic strategies

### Co-transferred Type 17 T cells are associated with resistance to TIL therapy

Having previously identified substantial *ex vivo* expansion and infiltration of Th17 in post-ACT lesions (Fig. 4b, g), we next investigated their contribution to TIL therapy outcomes.

Considering both CD4⁺ Th17 and CD8⁺ Tc17 cells, we observed an increased fraction of Type 17 T cells in TIL products of NRs (Fig. 7a). Subclustering of CD4⁺ and CD8⁺ T cells from TIL products of six patients similarly revealed enrichment of Th17 and Tc17 cells in NRs compared to Rs, suggesting a potential link between Type 17 T cell abundance and resistance to TIL therapy (Fig. 7b-e and Extended Data Fig. 7a, b). Given the small number of Rs in our cohort, we reanalyzed TIL products from other trials using comparable expansion protocols to validate our findings. DGE analysis of the cohort published by Chiffelle *et al.*^20^ similarly showed that TIL products from NRs exhibited increased expression of Type 17 T cell markers (*RORC*, *KLRB1, AQP3*, *IL26)* and other Th17/Tc17-associated genes (Fig. 7f and Extended Data Fig. 7e). Because subclustering did not resolve Type 17 T cells in this dataset, we applied gene signatures derived from our cohort, corroborating our finding that elevated Type 17 T cell numbers in TIL products are associated with non-response (Fig. 7g, and Extended Data Fig. 7f). Similarly, reanalysis of bulk RNA-seq data from Thompson *et al.*^33^ showed that CD4^+^ TIL products from NRs were enriched in genes associated with activated Type 17 T cells, supporting a link between Type 17 T cells and non-response (Fig. 7h, and Extended Data Fig. 7i).

**Fig. 7:**
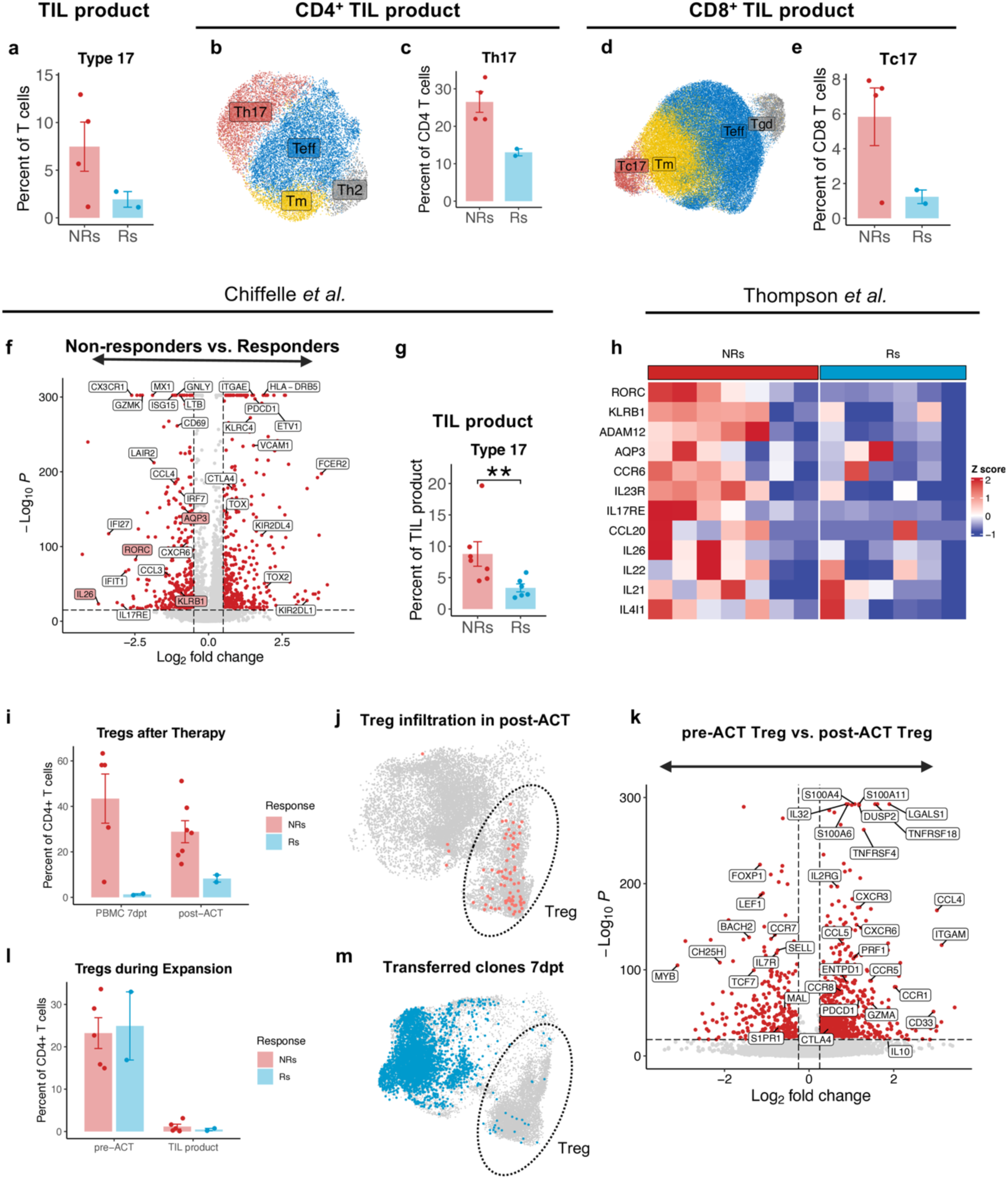
Co-transfer of Type 17 T cells and Treg expansion post transfer is associated with resistance to TIL therapy. **a)** Frequency of Type 17 cells in TIL products comparing NRs and Rs. Error bars indicate SEM. **b)** UMAP visualization of CD4^+^ TIL product. Tm, memory T cells; Teff, activated effector T cells; Th2, T helper 2 cells; Th17, T helper 17 cells. **c)** Frequency of Th17 cells in CD4^+^ TIL products. Error bars indicate SEM. **d)** UMAP visualization of CD8^+^ TIL products. Tm, memory T cells; Teff, activated effector T cells; Tc17, Type 17 CD8^+^ T cells; Tgd, gamma/delta T cells. **e)** Frequency of Tc17 cells in CD8^+^ TIL products. Error bars indicate SEM. **f)** DGE comparing TIL products of NRs (n=7) and Rs (n=6) on dataset by Chiffelle *et al.*^20^. **g)** Mean frequency of Type 17 T cells in the TIL product in the dataset by Chiffelle *et al*.^20^. Error bars indicate SEM. Statistical significance was assessed using Wilcoxon rank sum test. * p ≤ 0.05; ** p≤0.01; *** p≤0.001. **h)** Heatmap showing the expression of selected marker genes expressed by Th17 and Tc17 in the dataset by Thompson *et al.*^33^ with seven NRs and six Rs. **i)** Bar graph showing mean frequencies of Tregs in PBMC 7dpt and post-ACT. Error bars indicate SEM. **j)** Clones of Tregs found in PBMC 7dpt projected onto the UMAP of CD4^+^ T cells in post-ACT lesions. **k)** Volcano plot showing genes differentially expressed in Tregs comparing pre- vs. post-ACT in NRs. **l)** Bar graph showing mean frequencies of Tregs in pre-ACT and TIL products. Error bars indicate SEM. **m)** Clones of TIL drug product projected onto the UMAP in PBMC 7dpt highlighting overlap.

Overall, our results indicate an association between co-transferred Type 17 T cells and resistance to TIL therapy in NRs, validated across three independent datasets. Moreover, we found that Type 17 T cells are already present in baseline tumors and that TIL expansion protocols may inevitably promote their accumulation.

### NRs experience *de novo* expansion of peripheral Tregs after TIL transfer

Based on our previous observation of increased Treg accumulation in peripheral blood after TIL transfer, despite their low numbers in the TIL products, we next investigated their potential contribution to therapy outcomes (Fig. 4b). Notably, the accumulation of Tregs in PBMC 7dpt was only observed in NRs (Fig. 7i). Similarly, post-ACT lesions from NRs contained higher proportions of Tregs, whereas responding lesions showed reduced fractions (Fig. 7i). Clonal tracking further confirmed that peripheral Treg clones detected at 7dpt infiltrated post-ACT lesions in NRs, potentially influencing anti-tumor response (Fig. 7j). To further characterize these newly infiltrated Tregs, we performed DGE analysis comparing pre- and post-ACT Tregs of NRs. Post-ACT Tregs in NRs upregulated chemokine receptors (*CXCR3, CXCR6, CCR5, CCR1, CCR8*) and activation markers (*TNFRSF18, TNFRSF4, PDCD1, ENTPD1*), indicating active recruitment, migration, and immunosuppressive activity within the TME (Fig. 7k).

Further analysis suggested that pre-ACT lesions in NRs and Rs contained similar Treg proportions, and *ex vivo* expansion did not increase their numbers, indicating differential responses between Rs and NRs after TIL transfer (Fig. 7l).

As Tregs were nearly undetectable in the final TIL products, we further investigated their origin to rule out differentiation of other subpopulations. TCR clonal overlap analysis between the TIL products and PBMCs at 7 dpt revealed that these Treg clones did not originate from the transferred TILs, but instead expanded *de novo* in patients (Fig. 7m).

These data suggest that in NRs, early immunosuppression emerges post-transfer, characterized by increased Treg accumulation in peripheral blood. Absent in the TIL products, these Tregs expanded *de novo*, exhibiting features associated with tumor homing, likely contributing to an unfavorable TME that diminished therapeutic efficacy.

## Discussion

TIL-ACT has emerged as a promising approach for treating advanced melanoma, yet underlying T cell fate decisions during *ex vivo* expansion and resulting cellular composition of TIL products remain incompletely understood. In this study, we provide a comprehensive analysis of T cell dynamics during TIL therapy, with a focus on tumor-reactive subpopulations. By leveraging longitudinal patient samples and complementary datasets, we provide comprehensive characterization of tumor-reactive subpopulations during TIL expansion and after administration, highlight potential reasons for therapy resistance, and offer actionable insights to improve TIL-based therapies.

Consistent with our findings, the study by Chiffelle *et al.* found that during TIL expansion, tumor-reactive CD8^+^ TILs can be reinvigorated, downregulating tumor-associated transcriptional signatures, such as exhaustion markers^20^. However, in post-ACT lesions, these clones exhibited an exhausted effector state in Rs, whereas NRs showed poor infiltration of tumor-reactive CD8^+^ TILs, with low cytotoxic activity and low exhaustion marker expression^20^. Building on these insights, our results confirm plasticity of tumor-reactive CD8^+^ T cells during TIL therapy and uncover additional transcriptional and clonal dynamics. At baseline, these clones predominantly exhibited an exhausted phenotype, consistent with other previous studies^31,34^. During the first phase of *ex vivo* expansion (preREP), tumor-reactive clones preferentially expanded in response to IL-2. However, their growth plateaued or contracted in the second expansion phase (REP), likely due to polyclonal stimulation favoring bystander T cells. Notably, during preREP expansion, exhausted tumor-reactive CD8^+^ T cells acquired unique effector profiles characterized by high HLA-II expression, or, in the absence of recent TCR engagement, migratory profiles marked by high expression of *KLF2*. In contrast, the REP drove tumor-reactive CD8^+^ T cells towards a phenotype with increased cytotoxic and tumor homing potential. Post-transfer, tumor-reactive CD8^+^ clones in the peripheral blood exhibited stem-like profiles and expressed markers associated with lymph node egress resembling those identified by Tsui *et al.*^25^, suggesting both tumor homing capacity and the potential to establish a stem-like reservoir. After infiltration into post-ACT lesions, however, these T cells reacquired expression patterns associated with both anti-tumor activity and exhaustion. Importantly, differences emerged between Rs and NRs: in Rs, tumor-reactive T cells retained earlier developmental states with stem-like features, whereas in non-responders they shifted towards a terminally differentiated, dysfunctional phenotype with elevated expression of exhaustion markers. These findings differ to the observation by Chiffelle *et al.*, but align with previous studies linking stem-like CD8^+^ T cells to response to ICB^29^ and underscore the importance of maintaining stem-like T cell states for optimal TIL-ACT efficacy. While these findings highlight the dynamics of tumor-reactive CD8⁺ T cells during TIL therapy, tumor-responsive CD4⁺ T cells displayed distinct patterns that warranted separate investigation. In baseline tumors: two major subsets were observed: Th1 with their exhausted (Tex) counterpart and Tfh-like clones, latter associated with tertiary lymphoid structure formation and favorable clinical outcomes across multiple solid tumors^35^. Both subsets expanded during preREP; while Tex remained exhausted, Tfh were reinvigorated to an HLA-II-expressing state resembling that of tumor-reactive CD8^+^ T cells. In the REP phase, tumor-responsive CD4^+^ T clones especially Tex and Tfh subpopulations with strong anti-tumor potential, were progressively outgrown by other T cells, underscoring a critical opportunity for improving TIL therapy. Yet, they acquired Teff characteristics, including cytotoxic and tumor-homing markers. Post-transfer, these clones reappeared at lower frequencies in peripheral blood but in an activated stem-like state, expressing cytokines and migration markers suggesting active contribution to anti-tumor response. While tumor-reactive CD8^+^ clones were found exhausted and dysfunctional in post-ACT lesions of NRs, tumor-responsive CD4^+^ Tex shifted to more matured phenotypes with high cytotoxic potential. Notably, Tfh subpopulations also adopted a Th1-like phenotype characterized by elevated cytotoxic marker expression, suggesting TME remodeling potential despite ongoing exhaustion. Together, these findings indicate that tumor-responsive CD4^+^ T cell subsets may represent an underappreciated component of TIL therapy, warranting further investigation to improve therapeutic efficacy. Additionally, we identified distinct signatures for tumor-reactive CD8^+^ and tumor-responsive CD4^+^ subpopulations within the intermediate product (after preREP). These signatures offer opportunities to prospectively identify and isolate relevant clones during *ex vivo* expansion. Enriching for tumor-reactive clones at this stage may reduce bystander T cell contamination improving overall TIL potency. Current phenotypic enrichment strategies from tumors remain limited: Although markers such as PD-1, CD39, and TIGIT are associated with neoantigen reactivity, these subsets often fail to expand efficiently *in vitro*^36^. Furthermore, early dysfunctional tumor-reactive T cells, despite their greater proliferative potential, often lack robust expression of these canonical markers^37^. By contrast, during preREP we observed marked expansion of tumor-reactive subpopulations with distinct expression profiles, including *HLA-II*, *CD74*, *PDCD1*, *S1PR1*, *CD28*, and *CD70*. We propose leveraging these markers as enrichment markers in intermediate products, thereby enabling scalable isolation of tumor-reactive clones through approaches such as magnetic bead-based selection^38^.

Beyond tumor-reactive T cell dynamics, our study suggests that Type 17 cells, not showing anti-tumor reactivity in our assays, were preferentially enriched in TIL products from NRs. This association was further supported by two independent publicly available datasets, in which Type 17 T abundance also correlated with non-response. Although Type 17 T cells can exert anti-tumor effects when their TCR recognizes tumor antigen^39^, they are also capable of promoting tumor progression by systemic IL-17 production. IL-17 induces pro-survival and pro-angiogenic programs via IL-6-STAT3 signaling^40^ and promotes recruitment and expansion of immunosuppressive cells^41,42^. Furthermore, Th17 cells directly promote Tregs activation, expansion, and stability via TNFR2 stimulation^43–46^. Consistent with our findings, Knochelmann *et al.* showed that long-term expanded Th17 cells reduced therapeutic efficacy in mice by increasing Treg tumor infiltration^47^. Together, these findings provide evidence that co-transferred Type 17 T cells could undermine ACT efficacy by reinforcing immunosuppressive circuits. We therefore, knowing that treatment regimens in the analyzed trials differ, propose assessing Type 17 T cell abundance after *ex vivo* expansion as a potential prognostic biomarker for therapy response in larger patient cohorts.

Finally, our study underscores a potential role for Tregs in treatment resistance. Although IL-2 administration after TIL infusion is critical for engraftment and proliferation, it also was shown to promote Treg expansion especially using low dose IL-2 as used in our trial^48,49^. Also, in high dose IL-2 regimes peripheral Treg accumulation following TIL transfer has previously been correlated with non-response^50^, suggesting a potential resistance mechanism across different treatment modalities. In our cohort, peripheral Treg expansion was observed specifically in NRs. These Tregs did not originate from the infused TIL product but appear to emerge *de novo* post-transfer, exhibiting a strong immunosuppressive transcriptional profile. A link between the transfer of increasingly high numbers of activated Type 17 T cells in TIL product of NRs and the accumulation of peripheral Treg 7dpt cannot be ruled out and potential resistance mechanism remain to be elucidated. Peripheral Tregs were enriched in post-ACT lesions of NRs, where they may have promoted impaired effector function and heightened exhaustion of tumor-reactive T cells. Targeting this mechanism could enhance TIL therapy efficacy. For instance, ANV419, an antibody-IL-2 fusion protein that selectively promotes the expansion of effector T cells while limiting Treg expansion^51^, is currently evaluated in a phase I clinical trial (NCT05869539) and may offer a promising alternative to conventional IL-2 therapy for supporting TIL engraftment.

Despite the valuable insights gained in this study, the relatively small patient cohort and inter-patient variability remain limitations for correlating clinical outcomes, underscoring the need for validation in larger clinical trials. Furthermore, despite the use of largely standardized expansion protocols in our trail allows comparability, differences in treatment regimens after transfer, such as IL-2 dosing strategies or the addition of PD-1 blockade, may also influence outcomes and should be carefully considered by comparing with other studies.

Beyond these limitations, our findings advance the understanding of T cell plasticity in TIL therapy and highlight strategies to improve its therapeutic efficacy. Targeting tumor-reactive subpopulations, monitoring of Type 17 T cell abundance, and limiting Treg expansion represent actionable approaches to improve TIL therapy and overcome resistance. Incorporating these insights into next-generation expansion protocols and rational combination strategies may enable more consistent and durable responses in patients with metastatic melanoma.

## Materials and methods

### Patient population and trial design

The results of this clinical trial have been reported by König *et al.*^17^. Patients were eligible if they were 18 years or older and had histologically confirmed, unresectable stage III or IV cutaneous melanoma with disease progression after at least one anti-PD-1 based treatment line and an additional BRAF inhibitor in patients with BRAF V600 mutation, an accessible metastasis for tumor collection, measurable disease according to the Response Evaluation Criteria in Solid Tumors (RECIST), version 1.1, and Eastern Cooperative Oncology Group (ECOG) performance status score of 0 or 1 (on a 5-point scale, with higher scores indicating greater disability). The key exclusion criteria were non-cutaneous melanoma and active or untreated brain or leptomeningeal metastases.

In this academic phase I trial, patients underwent surgical excision of a metastasis for the generation of the TIL product. Nine patients were scheduled for TIL treatment. The TILs were manufactured at the GMP Facility for Advanced Therapies of the University Hospital of Basel. If the generation of the TIL product was successful, patients started lymphodepleting chemotherapy (LD) intravenously with cyclophosphamide at a dose of 60 mg/kg for days −7 and −6 and fludarabine daily at a dose of 25 mg/m^2^ (maximum of 50 mg) for days −5 to −1. Patients received TIL infusion on day 0, followed by daily subcutaneous administration of aldesleukin (interleukin-2, IL-2) at a dose of 125’000 IE/kg for 10 days, with a 2-day break after the 5^th^ dose. On day 14, patients were started on intravenous nivolumab at a dose of 240 mg and were thereafter treated with the same dose every 2 weeks. Treatment with nivolumab was continued until the occurrence of disease progression or for a maximum duration of 2 years. Patients received support with granulocyte colony-stimulating factor (G-CSF) after TIL infusion and further supportive treatment during TIL-ACT as needed. Prophylaxis for herpes simplex virus and pneumocystis was administered after TIL-ACT for 3 and 6 months, respectively.

The primary endpoint was the safety of the study intervention by assessing adverse events. Key secondary endpoints were the objective response rate (ORR), progression-free survival (PFS), and overall survival (OS). Tumor regression occurred in most patients (7/9) at the first imaging scan performed approximately one month after TIL-ACT. The best radiographic response according to RECIST v1.1 was partial response (PR) in 2 patients (ORR 22%) and stable disease (SD) in 3 patients. Four patients had progressive disease (PD) on the first imaging scan. All the patients experienced disease progression at one point. The median PFS after TIL transfer was 2.2 months. Five patients received subsequent systemic therapies, including the use of combined PD-1/CTLA-4 inhibitors in three patients. The median OS from TIL transfer was 7.2 months, with 5 and 3 patients alive 6 and 12 months after TIL transfer, respectively. For translational purposes, we classified two patients with PR as responders (Rs), and two patients with PD and three patients with SD as non-responders (NRs).

### Patient material processing

Resected solid tumor lesions were mechanically dissociated, digested with accutase (PAA Laboratories), collagenase IV (Worthington), hyaluronidase (MilliporeSigma), and DNAse type I (MilliporeSigma), filtered, washed, and frozen as single-cell suspensions for future use. Human PBMCs were isolated from whole blood by density gradient centrifugation using LymphoprepTM (Stemcell). PBMCs were cryopreserved for subsequent use. Aliquots of the TIL infusion products were frozen directly after harvest and stored in liquid nitrogen for later use. Excess intermediate products were obtained via the TIL expansion process.

### Fluorescence-activated cell sorting for single-cell RNA-sequencing

Cells were thawed, washed, and resuspended in staining buffer (PBS with 1% BSA) containing a Hu Fc Receptor Binding Inhibitor (Thermo Fisher Scientific). After 10 min of incubation at 4 °C, surface stain solution containing antibodies (CD3 BV421 Clone: SK7; CD45 PerCP-Cy5.5 Clone: 2D1) was added and incubated for 30 min at 4°C. The cells were washed twice and resuspended in staining buffer containing propidium iodide (1:1000; Thermo Fisher Scientific). Fluorescence-activated cell sorting (FACS) was performed using BD Melody or BD FACS Aria III. The cells were gated on single cells, live (PI-negative), CD45+, and CD3+. Cells were sorted in cold PBS with 2% FBS (Sigma-Aldrich) and used immediately for the single-cell immune profiling protocol (10x Genomics).

### Single-cell RNA and single-cell TCR sequencing

Single-cell TCR-seq and 5′ gene expression profiling of single-cell suspensions using the Chromium Single Cell V(D)J Reagent Kits v2 from 10x Genomics was performed according to the manufacturer’s instructions. Cells were loaded in a 10x Genomics cartridge to achieve a target recovery of approximately 10,000 cells for each sample. Cell-barcoded 5′ gene expression libraries and TCR libraries were sequenced using the Illumina NovaSeq6000 system by the Genomics Facility Basel. Gene expression and V(D)J reads were aligned in parallel and annotated using CellRanger v. 7.1. For this the human reference genome GRCh38 2020-A and human V(D)J reference genome GRCh38 ensembl-7.1.0. was used.

### Quality control and normalization of scRNA-seq data

The read count matrices were processed and analyzed using R v4.3.0, and Seurat v5.1.0, with default parameters for all functions unless otherwise specified^52^. Cells were filtered using nFeatures over 650, nCounts over 1250, mitochondrial gene percentage of less than 15%, ribosomal gene percentage of more than 5%, and hemoglobin gene percentage of less than 10%. Doublets were identified based on T cells containing more than one TCR-β chain and removed. Counts were normalized and regressed for cell cycle, percentage of mitochondrial genes, and interferon response genes^53^.

### Dimensional reduction and clustering

Genes irrelevant for clustering, for example, TCR genes, histone genes and ribosomal genes were excluded from variable features by using the gene blacklist used by Zheng *et al.*^19^. After PCA reduction, Harmony integration implemented in Seurat was used to integrate the dataset and remove patient- and sample-type-specific alterations. The 20 first Harmony corrected PCs were used to calculate the UMAP representation. For clustering the 20 first corrected PCs were used as input for the Leiden algorithm implemented in Seurat with resolutions from 0.3 to 0.6. T cell subsets were separated based on *CD8A, CD8B*, or *CD4* expression per cluster. The clustering procedure was repeated as described above until no CD8 or CD4 specific clusters were detected. The datasets were further purified from cells expressing *CD8A*, *CD8B*, or *CD4*. Cell type-specific clusters were identified using canonical marker genes and signatures, as published by Zheng *et al.*^19^. The R package UCell v2.6.2 was used for signature scoring. The R package Nebulosa v1.15.0 was used for gene expression visualization on UMAPs.

### scTCR-seq analysis

TCR count metrics were integrated into the Seurat object using scRepertoire v1.7.2. For quality control and data exploration purposes, the R packages scRepertoire v1.7.2, immunarch^54^, and scirpy v0.13.1 were used. For clone tracking, clonotypes were defined based on the nucleotide sequence of the CDR3-β region. The expansion score was defined as the log2-fold change in the frequency at which a clone was detected in two subsequent samples.

### Differential gene expression

DGE was performed using the in Seurat implemented function FindMarkers with a log fold change threshold set to zero. For patient-averaged differential gene expression of tumor-reactive vs. bystander T cells, an aggregated count matrix was created, and the standard DESeq2 v1.42.1 workflow was followed using sample origin as a covariate.

### RNA velocity trajectory analysis

Spliced and unspliced reads were counted using the velocyto.py package v0.17.17 from aligned bam files generated by CellRanger v7.1.0. TCR, histone, and ribosomal genes were excluded using the gene blacklist by Zheng *et al.*^19^. From the resulting count matrices top 2000 variable genes with a minimum of 20 shared counts were selected and normalized. The moments were calculated for velocity estimation and used in the dynamic model of scVelo to learn the unspliced/spliced phase trajectory. For visualization, the resulting trajectory vectors were embedded into the UMAP space.

### Pseudotime trajectory analysis

For pseudo temporal analysis we used the R package Monocle 3 v1.3.4. First, we loaded the pre-processed Seurat object into Monocle. After size factor estimation and combining all cells into one partition, we used the learn_graph function with default parameters to learn the cell connectivity. Next, we ordered the cells using the naïve T cell cluster Tn as root cells. We used spatial autocorrelation analysis implemented in Monocle 3 to infer differentially expressed genes along the pseudotime axis.

### Expansion of TCL

Autologous short-term cultured tumor cell lines (TCLs) were generated separately from TILs, either from tumor digests, tumor fragments, or after mincing, as previously described^55–57^. Adherent cells were passaged in RPMI 1640 medium supplemented with 10% FBS, 1 mM sodium pyruvate (Sigma-Aldrich), 100 U/mL penicillin, 100 µg/mL streptomycin (Thermo Fisher), and 500 ng/mL Solu-Cortef (Pfizer AG).

### Anti-tumor reactivity assay

To assess tumor-reactivity, the co-culture assay described by Andersen *et al.*^56^ was adapted. Briefly, TILs were thawed and rested for two days in assay medium (RPMI-1640 plus GlutaMAX with 1 mM sodium pyruvate (Sigma-Aldrich), 100 U/mL penicillin, 100 µg/mL streptomycin (Thermo Fisher), and 10% heat-inactivated AB Human serum). The TILs were washed, CFSE-labeled (Thermo Fisher Scientific), and co-cultured at 37°C for 10 h with fresh tumor digest (FTD) or autologous short-term cultured tumor cell lines (TCLs) at an effector/target (E/T) ratio of 1:1. The TCLs were pre-treated for 72 h with 100 IU/ml interferon-gamma (IFNγ; Imukin, Boehringer-Ingelheim). Monensin (BioLegend) and Brefeldin A (BioLegend) were added (dilution 1:1000 each) at the beginning of the co-culture. As negative controls, TCR-HLA interactions were blocked with anti-HLA-A, B, C (clone: W6/32, BioLegend, 20 µg/ml) and anti-HLA-DR, DP, DQ (clone: Tü39, BioLegend, 20 µg/ml).

### Fluorescence-activated cell sorting for anti-tumor reactivity assay

For intracellular staining procedures while preserving high RNA quality, the instructions of Healy *et al*.^58^ were followed. First, cells were washed with PBS and stained with Zombie NIR (BioLegend) for live/dead labeling. After washing in PBS with 1% FBS, the cells were fixed and permeabilized in a solution containing 4% PFA (Thermo Fisher Scientific), 0.1% saponin (Sigma-Aldrich), and 1 unit/μL of RNasin Plus RNase inhibitor (Promega) on ice for 20 min in the dark. Cells were then washed with permeabilization buffer (0.1% saponin, 2 M NaCl (Thermo Fisher Scientific), 0.5% BSA (Sigma-Aldrich) in RNase-free PBS (Thermo Fisher Scientific)), and intracellular staining was performed in permeabilization-stain buffer (PBS with 0.1% saponin containing 1 unit/μL of RNasin Plus RNase inhibitor) supplemented with anti-TNF⍺-APC (clone: MAB11, Biolegend) and anti-IFNɣ-BV421 (clone: 45.B3, Biolegend) at 4°C for 45 min. Intracellular staining was performed at a density of 10 × 10^6^ cells/mL. The stained cells were then washed twice with permeabilization buffer and resuspended in sorting buffer (1% BSA, 2 M NaCl in RNase-free PBS). TNF⍺ or IFNɣ positive live populations were sorted using BD SORPAria III.

### Total RNA isolation from sorted T cells

Cells were isolated from the sort buffer by centrifugation at 2000 RCF for 15 min at 4°C. Total RNA and DNA were extracted simultaneously using the AllPrep DNA/RNA FFPE Kit (Qiagen), with some modifications from the vendor’s instructions. Briefly, the deparaffinization step was skipped and directly followed by protein digestion, which was extended for 1 h at 56°C at 300 rpm. After the separation of total DNA and RNA, incubation at 80°C for partial reversion of formaldehyde modifications was not performed and directly followed by the addition of RLT buffer and ethanol. All samples were eluted with 16 µL pre-warmed RNase-free water to increase the final RNA concentration. Total RNA concentration and quality were determined using a high-sensitivity RNA screen type assay (Agilent) on a 4200 TapeStation System (Agilent).

### T cell receptor library preparation and sequencing

Immediately after total RNA isolation, cDNA was generated using the Ion Torrent NGS Reverse Transcription Kit (Thermo Fisher Scientific). The maximum RNA input was loaded for cDNA synthesis to capture a large pool of T-cell clones. Next-Generation Sequencing (NGS) libraries were prepared using the Oncomine TCR Beta-SR RNA Assay (Thermo Fisher Scientific), according to the manufacturer’s instructions. Amplified and barcode-ligated libraries were purified using AMPure XP Reagent (Beckman Coulter). The library quality check was performed using the High Sensitivity D1000 Screen Tape Assay (Agilent) on a 4200 TapeStation System (Agilent) and quantified using the Ion Universal Library Quantitation Kit (Thermo Fisher Scientific). The library pool was prepared by combining equal volumes of libraries at a concentration of 50 pmol/L and loaded into Ion 550™ Chip (Thermo Fisher Scientific). The libraries were sequenced using an Ion GeneStudio S5 Prime Sequencer (Thermo Fisher Scientific). FASTQ files were generated using IonTorrent Server (Thermo Fisher Scientific).

### TCR alignment and annotation of bulk TCR-seq

Read extraction and clonality counts were obtained from the FASTQ files using MiXCR v4.4.1 with the implemented presets for Oncomine™ TCR Beta-SR RNA Assay^59^. Count tables were organized and explored using Immunarch R package^54^. We determined the log2 counts per million (CPM) and log2-fold changes in the shared TCR sequences between the bulk TIL product and the positively sorted population. Only clones that were shared between both groups and positively enriched in the sorted populations were classified as tumor-reactive. Tumor reactivity was verified in pre- or post-ACT samples using NeoTCR signatures or co-expression of tumor-reactivity signatures, exhaustion markers, and *CXCL13*, as described in the literature, as specific for tumor-reactive T cells^31,34,60,61^.

### Statistical methods

The R package geom_signif was used to evaluate statistical significance^62^. Statistical tests and parameters used are indicated in the figure legends. Prism 10 was used for visualization and statistical testing of mouse data.

## Data availability

All original data is available from the corresponding author upon reasonable request. scRNA-seq and scTCR-seq data are available at the European Genome-Phenome Archive under the accession ID: EGAP50000000107.

## Code availability

The code generated and used for the analysis of sequencing data are available from the corresponding author on reasonable request.

## Authors contributions

M. T. Sandholzer, A. G. Liner, and S. Uzun performed experiments and analyzed data; H. Thut, R. Ritschard, A. Zingg., N. R. Mantuano performed GMP manufacturing of infusion products; H. Läubli and A. Zippelius provided supervision; M. T. Sandholzer, A. G. Liner, and H. Läubli contributed to experimental design; M. T. Sandholzer, C. Serger, A. G. Liner, and H. Läubli prepared the manuscript; H. Läubli, and A. Zippelius provided funding; H. Läubli, and A. Zippelius led the research program.

## Acknowledgments

We thank the patients for allowing us to report their clinical information and data. We thank Werner Krenger and Anke Wixmerten for the support of the GMP productions. In addition, we thank Julia Manzetti, Dimitrios Tsakiris, and Astrid Tschan-Plessl for their initial help during the setup phase of the TIL program at the University Hospital Basel. We thank Francesc Baixauli and Nadia Schlauri for their support and material for the Th17 polarization. We thank the Flow Cytometry Facility and Animal Facility of the Department of Biomedicine at the University Hospital Basel for their daily contributions. Additional thanks to the Genomics Facility Basel and Christian Beisel for sequencing of the single-cell RNA libraries. We thank Johanna Nimmerfroh and Jonas Fürst for their great help with the mouse experiments. We thank Petra Herzig for helping with the patient samples. This work was supported by funding from the Fond’Action Foundation (to H. Läubli), research support and drug supply of nivolumab by Bristol Myers Squibb, and internal funding of the University Hospital Basel (Innovation Focus Cell Therapy). Research in the L.T. Jeker lab is supported by the European Research Council under the European Union’s Horizon 2020 Research and Innovation Program (grant agreement No. 818806).

