## Supplementary Data for "Longitudinal profiling of tumor-reactive T cells during TIL therapy in metastatic melanoma"

### Extended Data

**Extended Data Table 1: Patient metadata table used for single cell experiment.**

| Patient | Response | Gender | Transfused cells | time to post-ACT.1 | time to post-ACT.2 |
|---|---|---|---|---|---|
| UPN001 | PR | male | 74.25 x 10e9 | 21 weeks | 62 weeks |
| UPN002 | PD | male | 50 x 10e9 | NA | NA |
| UPN003 | PR | female | 70 x 10e9 | NA | NA |
| UPN006 | SD | male | 57 x 10e9 | 6 weeks | 15 weeks |
| UPN008 | SD | female | 66.25 x 10e9 | 11 weeks | 33 weeks |
| UPN009 | SD | female | 68.25x 10e9 | 31 weeks | 31 weeks |
| UPN011 | PD | male | 63.25 x 10e9 | 8 weeks | NA |

**Extended Data Table 2: T cell count per sample after filtering.**

| Patient | pre-ACT | Intermediate product | TIL product | PBMC 7dpt | post-ACT.1 | post-ACT.2 | PBMC_Ctrl |
|---|---|---|---|---|---|---|---|
| UPN000 | NA | NA | NA | NA | NA | NA | 13176 |
| UPN001 | 3184 | 1864 | 6396 | 10593 | 6114 | 2232 | NA |
| UPN002 | 2806 | 7590 | 9034 | 11379 | NA | NA | NA |
| UPN003 | 5796 | 8401 | 8008 | 17328 | NA | NA | NA |
| UPN006 | 12466 | 8222 | 11289 | 14909 | 424 | 331 | NA |
| UPN008 | 28878 | 12572 | 14562 | 16972 | 7128 | 5627 | NA |
| UPN009 | 9824 | 6352 | 12977 | 10684 | 13644 | 8046 | NA |
| UPN011 | 6149 | 10932 | 18634 | 2305 | 6900 | NA | NA |

**Extended Data Table 3: T cell count with identified TCRβ CDR3 regions per sample after filtering.**

| Patient | pre-ACT | Intermediate product | TIL product | PBMC 7dpt | post-ACT.1 | post-ACT.2 | PBMC_Ctrl |
|---|---|---|---|---|---|---|---|
| UPN000 | NA | NA | NA | NA | NA | NA | 11981 |
| UPN001 | 2790 | 1491 | 5725 | 9084 | 5208 | 1615 | NA |
| UPN002 | 1648 | 2053 | 1504 | 4718 | NA | NA | NA |
| UPN003 | 5086 | 6531 | 5926 | 14394 | NA | NA | NA |
| UPN006 | 11832 | 7234 | 10189 | 13255 | 391 | 287 | NA |
| UPN008 | 26113 | 10101 | 12094 | 13100 | 6531 | 4428 | NA |
| UPN009 | 8302 | 4826 | 10482 | 9474 | 10128 | 6161 | NA |
| UPN011 | 4613 | 7653 | 15422 | 2087 | 5936 | NA | NA |

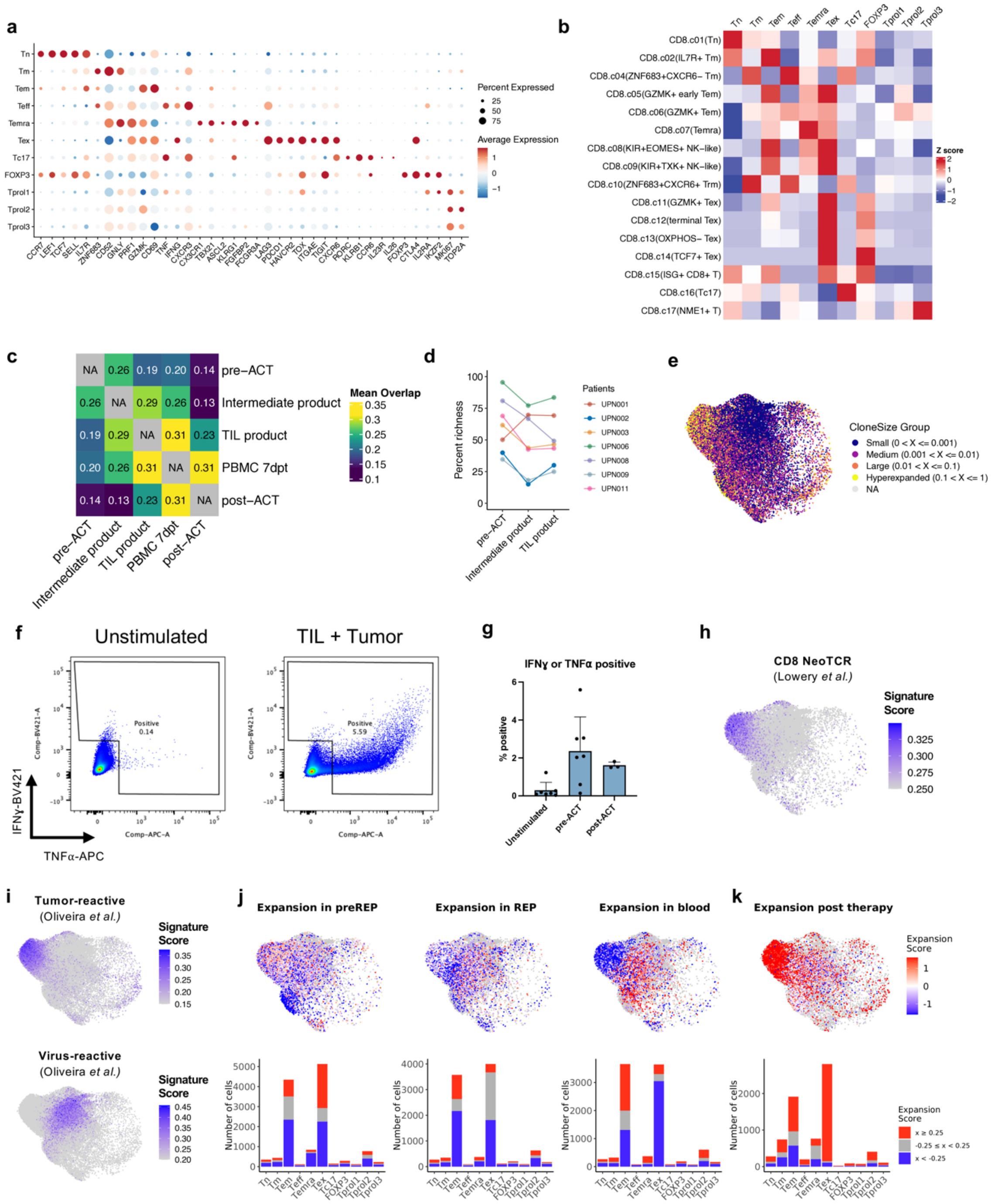


**Extended Data Fig. 1: Plasticity and clonal dynamics of tumor-reactive CD8$^+$ T cells. a)** Dot plot showing the expression of selected marker genes used for cluster annotation. **b)** Heatmap depicting the signature mapping used for cluster annotation. Signatures were derived from Zheng *et al.*[1] **c)** Heatmap depicting mean TCR overlap of CD8$^+$ T cells between samples. **d)** Clonal diversity as percent richness between pre- and post-ACT. **e)** Clonal expansion highlighted on UMAP. **f)** Example gating used for sorting of TNFα or IFNɣ positive populations. **g)** Bar plot indicating TNFα or IFNɣ positive fractions per condition. Mean ± SD. **h)** Signature score of neoantigen-reactive CD8$^+$ T cells by Lowery *et al.*[2] projected onto UMAP of tumor derived T cells. **i)** Signature score of tumor-reactive and virus-reactive T cells by Oliveira *et al.*[3] projected onto UMAP of tumor derived T cells. **j)** Expansion score highlighted in the UMAP representation of pre-ACT (top row). The number of cells per cluster classified as expanding (expansion score ≥ 0.25, red), contracting (expansion score < -0.25, blue), or unchanged (-0.25 ≤ x < 0.25, gray) (bottom row). **k)** Expansion score highlighted in the UMAP representation of post-ACT (top row). The number of cells per cluster classified as expanding (expansion score ≥ 0.25, red), contracting (expansion score < -0.25, blue), or unchanged (-0.25 ≤ x < 0.25, gray) (bottom row).

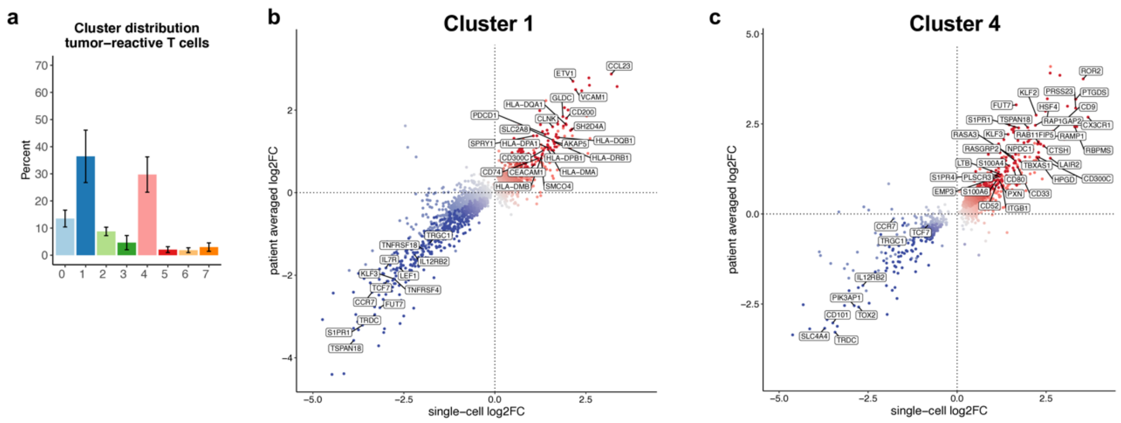


**Extended Data Fig. 2: Tumor-reactive CD8$^{+}$ T cells acquire distinct expression profiles during reinvigoration. a)** Cluster distribution of tumor-reactive T cells in clusters of intermediate products. **b-c)** Differential gene expression comparing tumor-reactive CD8$^{+}$ subpopulations in corresponding cluster with remaining T cells in intermediate products using patient bulked comparison and single-cell comparison. Genes significantly upregulated in both comparisons were highlighted in dark color and genes significantly upregulated only in single-cell comparison in light color.

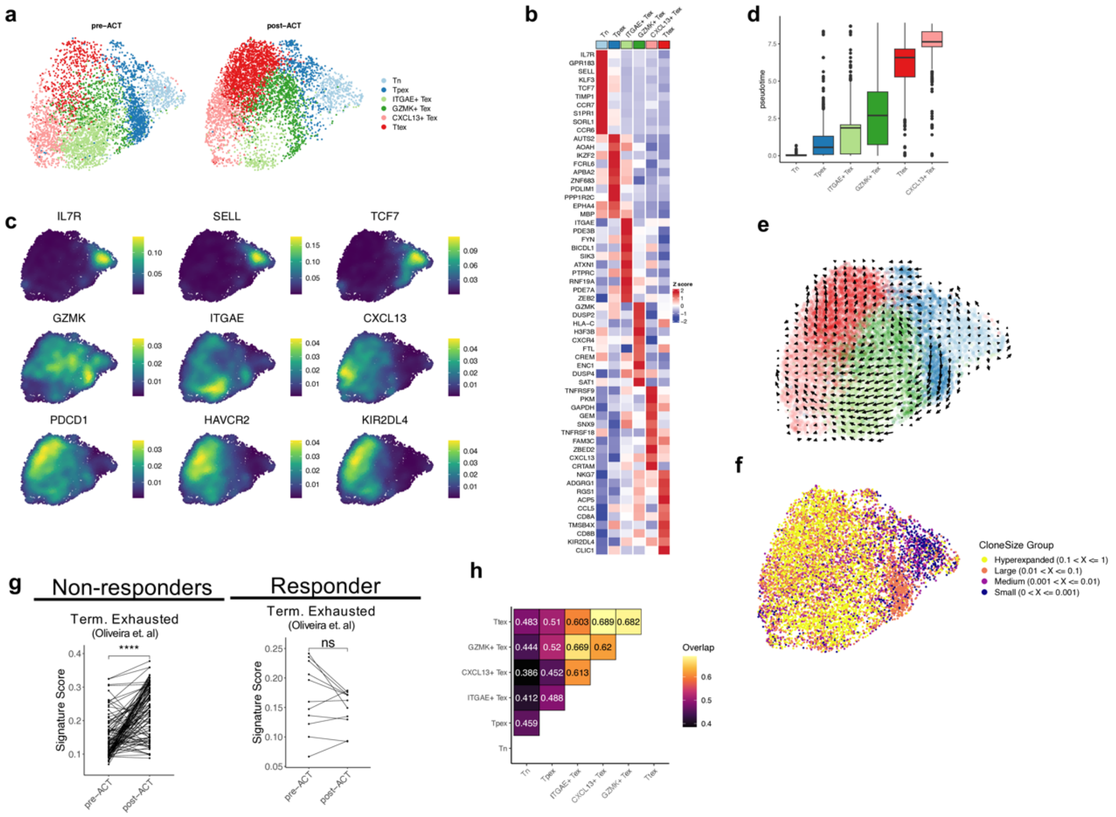


**Extended Data Fig. 3: CD8$^{+}$ tumor-reactive T cells show an elevated dysfunctional profile in post-ACT lesions of non-responders. a)** UMAP visualization of the cell states split into pre- and post-ACT lesions. **b)** Heatmap showing the top genes differentially expressed between clusters. **c)** UMAP plots showing the expression levels of selected marker genes. **d)** Pseudo-time values per cell grouped by clusters. **e)** RNA velocities calculated based on unspliced vs. spliced RNA embedded in UMAP. **f)** Expansion state highlighted on UMAP representation. **g)** Signature scores of published signatures comparing mapped clones detected in pre- and post-ACT lesions of Rs and NRs[3]. Statistical significance was assessed using Wilcoxon rank sum test. * p ≤ 0.05; ** p≤0.01; *** p≤0.001; **** p≤0.0001 **h)** Heatmap depicting clonal overlap between clusters.

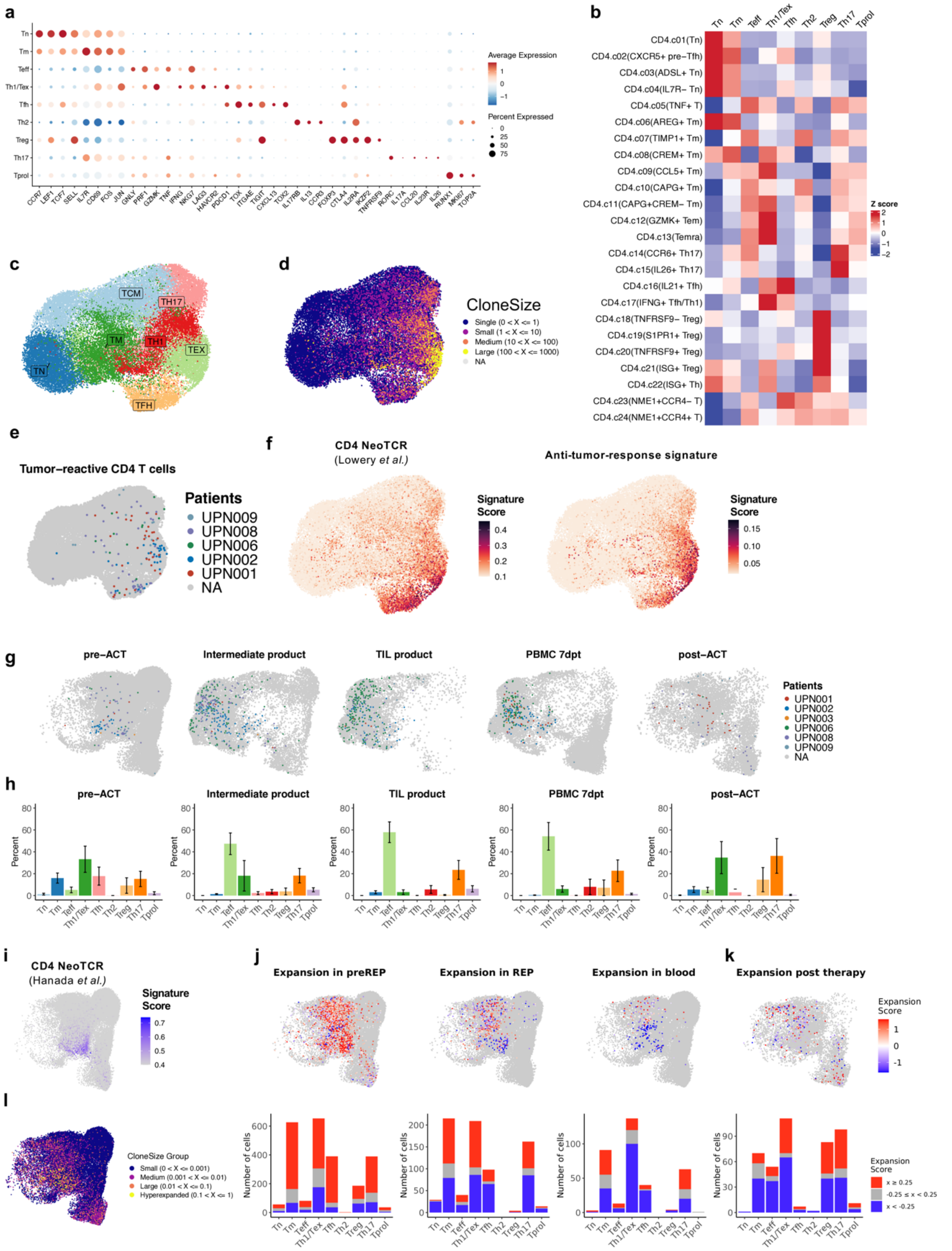


**Extended Data Fig. 4: Plasticity and clonal dynamics of tumor-responsive CD4$^{+}$ T cells. a)** Dot plot showing the expression of selected marker genes used for cluster annotation. **b)** Heatmap depicting the signature mapping used for cluster annotation. Signatures were derived from Zheng *et al.*[1]. **c)** Subclustering of conventional CD4$^{+}$ T cells of pre-ACT and post-ACT samples. TN, naïve T cells; TCM, central memory

T cells; TM, memory T cells; TH1, T helper 1 cells; TH17 T helper 17 cells; TFH, follicular helper T cells; TEX, exhausted T cells. **d)** Clonal expansion highlighted on subclustering of conventional CD4$^+$ T cells. **e)** Tumor-reactive CD4$^+$ T cells highlighted on subclustering of conventional CD4$^+$ T cells. **f)** CD4 Neoantigen-reactive T cell signature by Lowery *et al.*[2] and Anti-tumor-response signature created in this study highlighted on UMAP. **g)** Tumor-reactive CD4$^+$ T cells highlighted in the UMAP representation at collected time points. **h)** Bar plots showing the mean CD4$^+$ T cell composition of the tumor-reactive subpopulation per time point. Error bars indicate SEM. **i)** Signature score of the CD4$^+$ neoantigen-reactive T cell signature by Hanada *et al.*[4]. **j)** Expansion score highlighted in the UMAP representation of pre-ACT (top row). The number of cells per cluster classified as expanding (expansion score ≥ 0.25, red), contracting (expansion score < -0.25, blue), or unchanged (-0.25 ≤ x < 0.25, gray) (bottom row). **k)** Expansion score highlighted in the UMAP representation of post-ACT (top row). The number of cells per cluster classified as expanding (expansion score ≥ 0.25, red), contracting (expansion score < -0.25, blue), or unchanged (-0.25 ≤ x < 0.25, gray) (bottom row). **l)** Clonal expansion highlighted in UMAP of pre-ACT CD4$^+$ T cells.

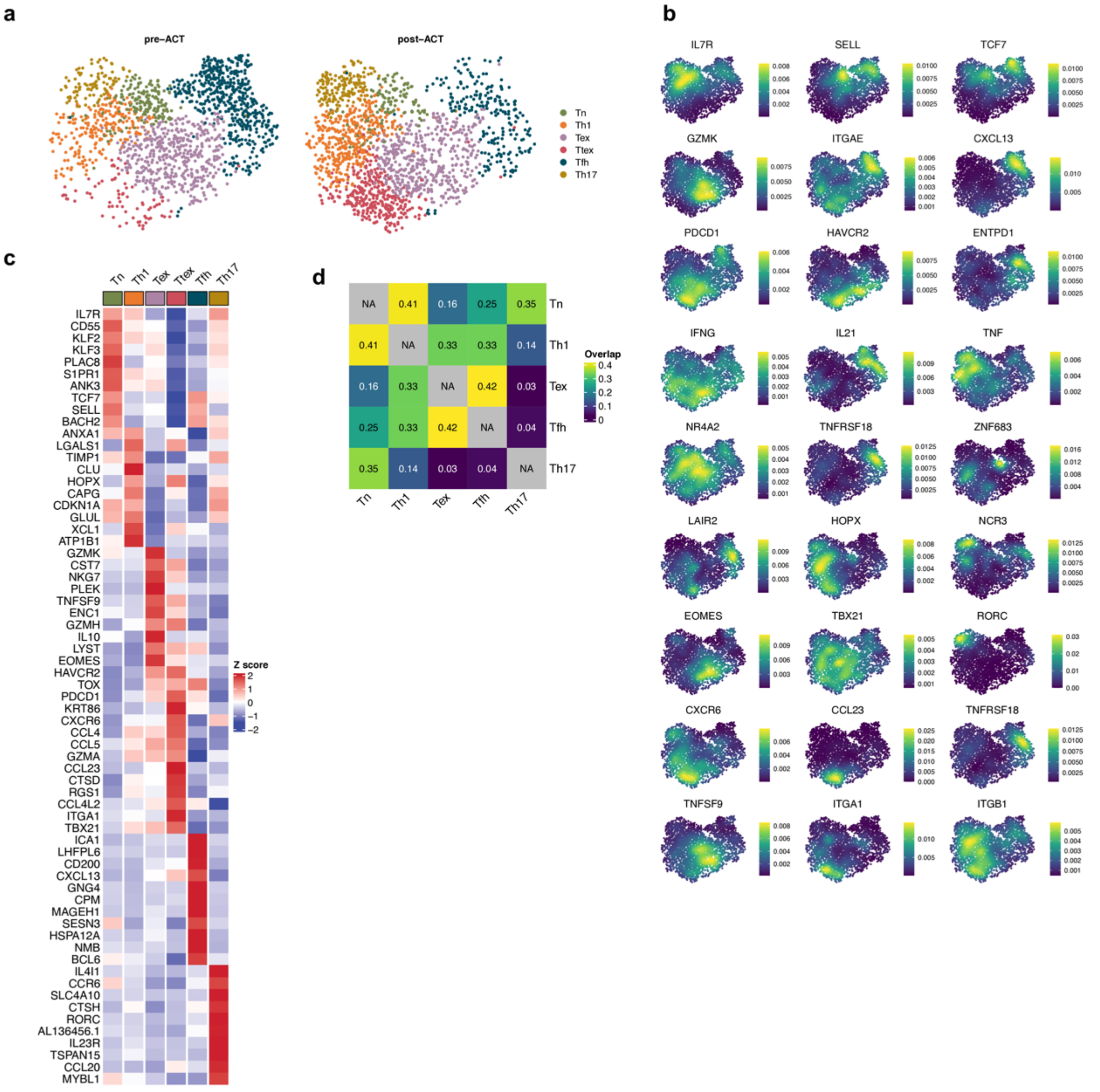


**Extended Data Fig. 5: Tumor-responsive Tfh and Tex retain effector function despite immunosuppressive TME. a)** UMAP visualization of tumor-responsive CD4$^+$ T cell states split into pre- and post-ACT lesions. **b)** UMAP plots showing the expression levels of selected marker genes. **c)** Heatmap showing the top genes differentially expressed between clusters **d)** Heatmap presenting clonal overlap between clusters.

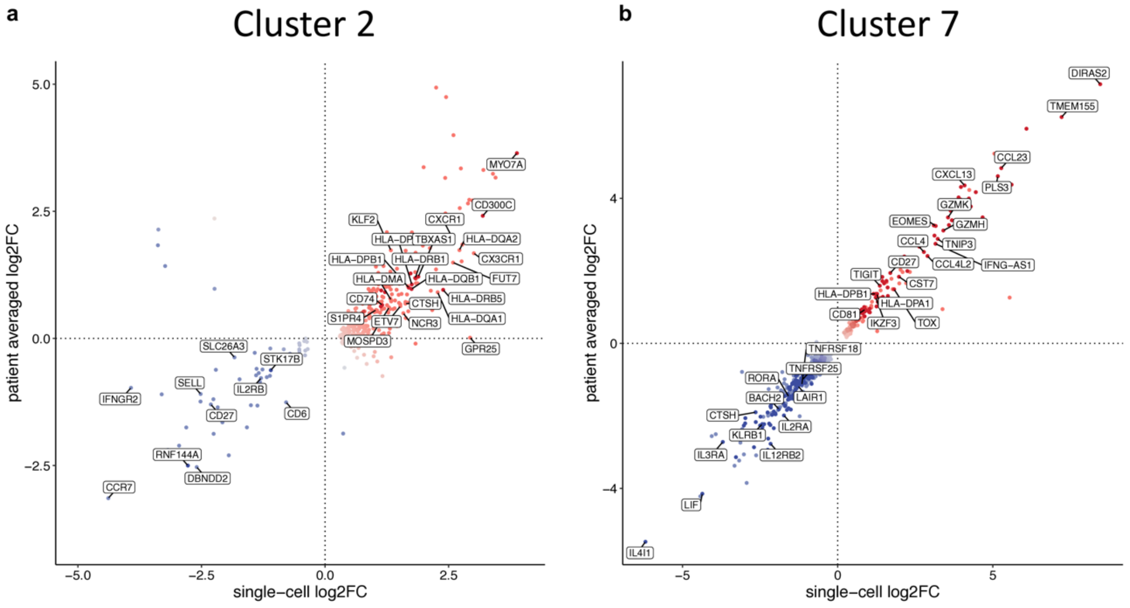


**Extended Data Fig. 5: Tumor-responsive Tfh but not Tex were efficiently reinvigorated during preREP. a-b)** Differential gene expression comparing tumor-responsive CD4$^{+}$ subpopulations in corresponding cluster with remaining T cells in intermediate products using patient bulked comparison and single-cell comparison. Genes significantly upregulated in both comparisons were highlighted in dark color and genes significantly upregulated only in single-cell comparison in light color.

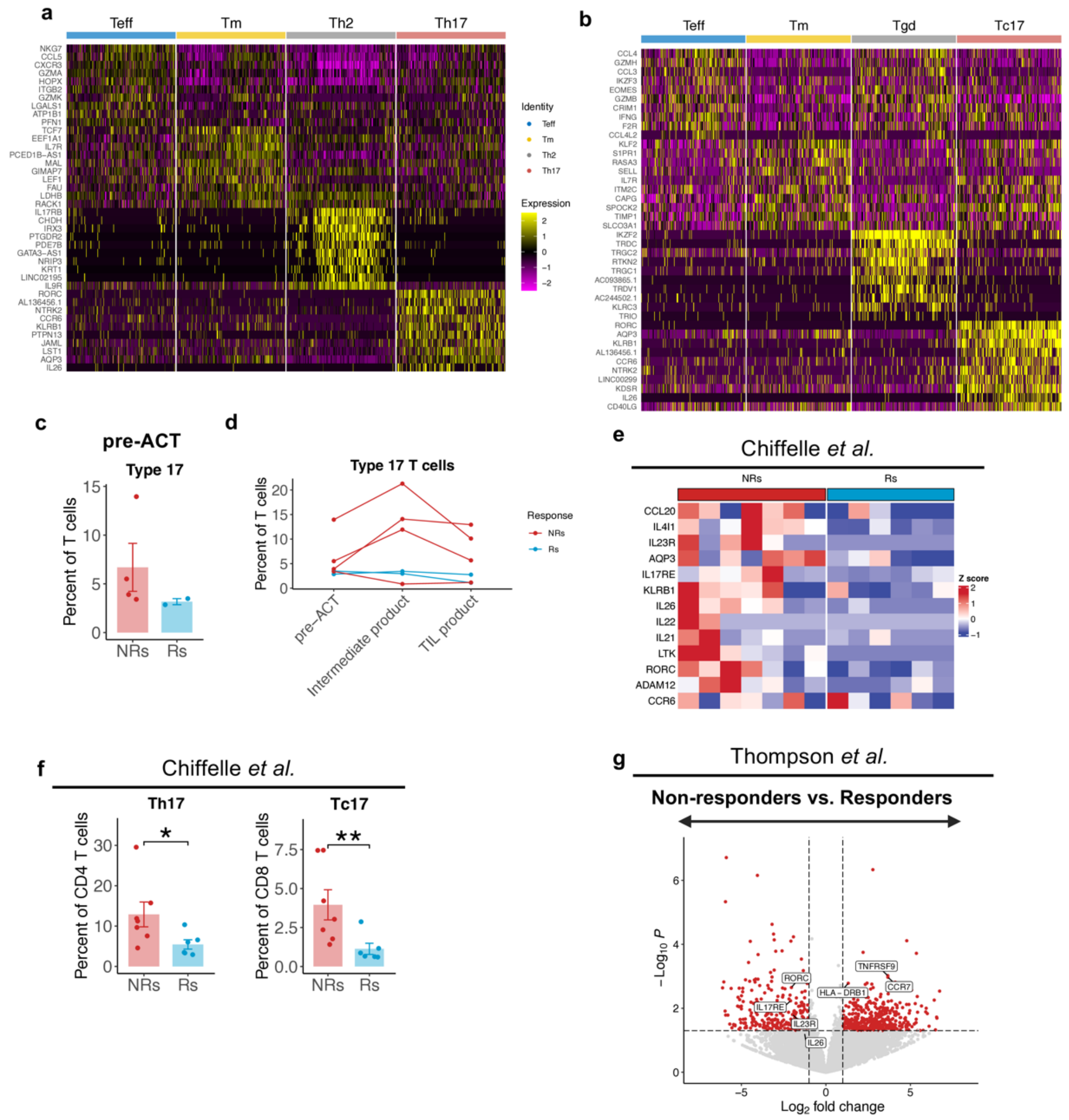


**Extended Data Fig. 7: Co-transfer of Type 17 T cells and Treg expansion post transfer is associated with resistance to TIL therapy. a)** Heatmap showing the top genes differentially expressed between clusters of CD4 TIL products. **b)** Heatmap showing top genes differentially expressed between clusters of CD8 TIL product. **c)** Mean frequency of Type 17 T cells in pre-ACT lesions in the dataset by Chiffelle *et al.*[5]. Error bars indicate SEM. Statistical significance was assessed using Wilcoxon rank sum test. **d)** Frequency change of Type 17 cells during TIL expansion. **e)** Heatmap showing the expression of selected Th17 and Tc17 marker genes expressed in the dataset by Chiffelle *et al.*[5]. **f)** Mean frequency of Th17 cells in CD4 TIL and Tc17 cells in CD8 TIL products in the dataset by Chiffelle *et al.*[5]. Error bars indicate SEM. Statistical significance was assessed using Wilcoxon rank sum test. * $p \leq 0.05$; ** $p \leq 0.01$ **g)** DGE comparing CD4 TIL products of NRs and Rs on the dataset by Thompson *et al.*[6].
